# Integrin Mechanosensing relies on Pivot-clip Mechanism to Reinforce Cell Adhesion

**DOI:** 10.1101/2023.12.13.571593

**Authors:** Andre R. Montes, Anahi Barroso, Wei Wang, Grace D. O’Connell, Adrian B. Tepole, Mohammad R.K. Mofrad

## Abstract

Cells intricately sense mechanical forces from their surroundings, driving biophysical and biochemical activities. This mechanosensing phenomenon occurs at the cell-matrix interface, where mechanical forces resulting from cellular motion, such as migration or matrix stretching, are exchanged through surface receptors, primarily integrins, and their corresponding matrix ligands. A pivotal player in this interaction is the *α*_5_*β*_1_ integrin and fibronectin (FN) bond, known for its role in establishing cell adhesion sites for migration. However, upregulation of the *α*_5_*β*_1_-FN bond is associated with uncontrolled cell metastasis. This bond operates through catch bond dynamics, wherein the bond lifetime paradoxically increases with greater force. The mechanism sustaining the characteristic catch bond dynamics of *α*_5_*β*_1_-FN remains unclear. Leveraging molecular dynamics simulations, our approach unveils a pivot-clip mechanism. Two key binding sites on FN, namely the synergy site and the RGD (arg-gly-asp) motif, act as active points for structural changes in *α*_5_*β*_1_ integrin. Conformational adaptations at these sites are induced by a series of hydrogen bond formations and breaks at the synergy site. We disrupt these adaptations through a double mutation on FN, known to reduce cell adhesion. A whole-cell finite element model is employed to elucidate how the synergy site may promote dynamic *α*_5_*β*_1_-FN binding, resisting cell contraction. In summary, our study integrates molecular and cellular-level modeling to propose that FN’s synergy site reinforces cell adhesion through enhanced binding dynamics and a mechanosensitive pivot-clip mechanism. This work sheds light on the interplay between mechanical forces and cell-matrix interactions, contributing to our understanding of cellular behaviors in physiological and pathological contexts.

**SIGNIFICANCE:** *α*_5_*β*_1_ integrin serves as a mediator of cell-matrix adhesion and has garnered attention as a target for impeding cancer metastasis. Despite its importance, the mechanism underlying the formation of a catch bond between *α*_5_*β*_1_ integrin and its primary ligand, fibronectin, has remained elusive. Our study aims to address this gap by proposing a pivot-clip mechanism. This mechanism elucidates how *α*_5_*β*_1_ integrin and fibronectin collaboratively reinforce cell adhesion through conformational changes induced by the dynamic interaction of a key binding motif known as the synergy site.

## INTRODUCTION

Adhesion bonds enable cells to interact dynamically with their surrounding environment, orchestrating the regulation of essential cellular processes such as proliferation, differentiation, and apoptosis (1–5). Integrins are transmembrane, heterodimeric proteins that play an important role in cell adhesion by tethering the inside and outside of the cell via binding partners in the extracellular matrix (ECM) (6). *α*_5_*β*_1_ integrin is one of 24 integrin heterodimers present in mammals (4) and mediates cell-tissue homeostasis by binding to its primary ligand, fibronectin (FN) (7, 8). *α*_5_*β*_1_ and FN are linked together at the RGD (Arg-Gly-Asp) motif and stabilized by the eight-amino-acid-long DRVPHSRN synergy site on FN (9), allowing extracellular and cytoplasmic forces to be transmitted across the cell membrane. The accumulation of *α*_5_*β*_1_-FN bonds form the basis for nascent cell adhesion and cell motion. Beyond *α*_5_*β*_1_-FN’s role in maintaining cell-tissue homeostasis, it has been implicated as a potential therapeutic target for cancer (10–12). For example, dysfunctional and overexpressed integrin bonds are markers of uninhibited cancer cell migration (13, 14). As such, numerous antagonists have been developed to attenuate integrin bonds, aiming to impede the invasion of multiple cancer cell types. Despite considerable efforts, these antagonists have faced challenges, demonstrating limited success in effectively preventing cancer cell invasion. (15, 16). Therefore, a better understanding of the biophysical nature of the *α*_5_*β*_1_-FN bond is needed to reveal mechanisms that can be exploited to target metastasis.

*α*_5_*β*_1_ integrin creates a catch bond with FN (9, 17, 18), which is a type of bond that increases in lifetime with greater applied force. The *α*_5_*β*_1_-FN catch bond allows for strong adhesion at the leading edge of a migrating cell and a steady release of the bond at the cell’s trailing end. Catch bonds have inspired development of synthetic catch bonds for manufacturing resilient materials (19–21). However, the mechanisms involved in the *α*_5_*β*_1_-FN catch bond’s ability to maintain its characteristic strength is unknown. Understanding the underlying mechanism of *α*_5_*β*_1_-FN catch bond resilience could identify structural protein characteristics that can be targeted to arrest cancer cells through substrate or protein modifications. Moreover, structural dynamics that enable catch bond behavior may inspire development of resistant nanomaterials with self strengthening properties. Ideally, the *α*_5_*β*_1_-FN catch bond could be imaged while an applied force is applied with a single-molecule testing setup (e.g., optical trap or magnetic tweezers). However, current atomic-resolution molecular imaging techniques, like cryo-EM and x-ray crystallography, require immobilizing the protein, making visualization of *in situ* structural changes of *α*_5_*β*_1_-FN challenging. In light of these experimental limitations, molecular dynamics (MD) simulations have been used to visualize protein conformational changes over time (22, 23).

Given *α*_5_*β*_1_-FN’s critical role in mechanosensing via its elusive catch bond dynamics, we used MD simulations to visualize the motion of *α*_5_*β*_1_-FN when acted on by an external load. We introduce a “pivot-clip” mechanism to model the *α*_5_*β*_1_-FN’s catch bond-like behavior, where the RGD motif acts as a stable pivot for FN about *β*_1_ integrin and the synergy site acts as a reinforcing clip connecting FN to *α*_5_. Past experiments demonstrated that mutating the synergy site diminishes catch bond behavior and weakens whole-cell and single molecule adhesion to *α*_5_*β*_1_ (18, 24). Even so, a lack of the synergy site does not significantly limit cell traction on a 2D substrate under minimal contractility (25). To explain how the synergy site may promote *α*_5_*β*_1_-FN binding while maintaining cell traction, we developed a 2D finite element (FE) model of the adhesive interface. Based on our MD and FE models, we present a theory that the synergy site in FN reinforces cell adhesion via stronger binding affinity and a mechanosensitive pivot-clip mechanism.

## MATERIALS AND METHODS

### Steered Molecular Dynamics Simulations

Constant velocity, all-atom steered MD simulations of the ectoplasmic *α*_5_*β*_1_-FN complex were run in GROMACS 2020.4 (26). The 7NWL crystal structure file of the *α*_5_*β*_1_-FN complex with the TS2/16 Fv-clasp was downloaded from the protein data bank. The *α*_5_*β*_1_ integrin head domain and the FN type III fragment 7-10 were isolated using PyMOL (27). We used MODELLER 10.4 (28) to impose a virtual R1374/9A double mutation, switching the arginine residues in positions 1374 and 1379 in FN to alanine (Figure 1B).

**Figure 1.**
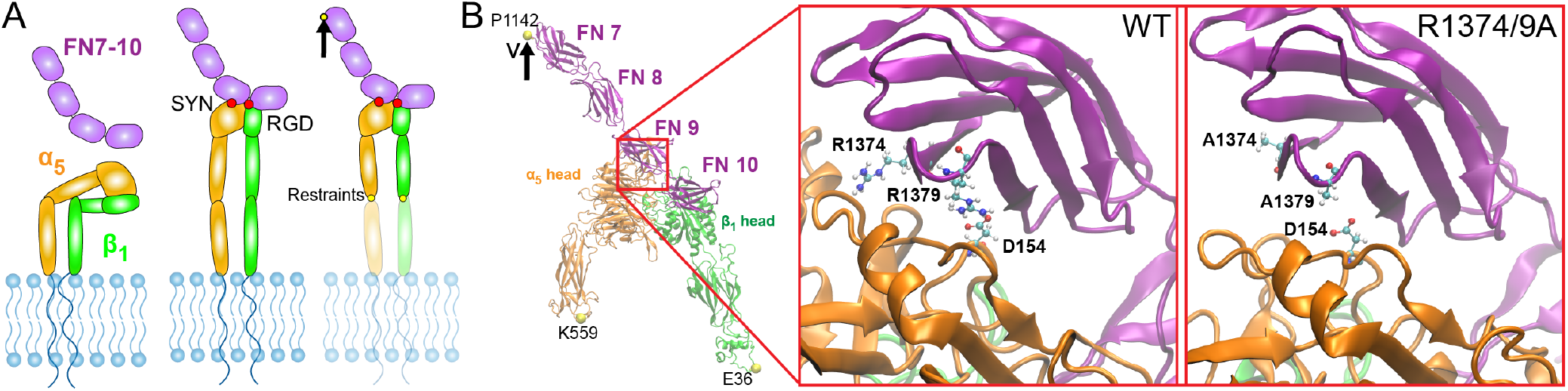
A) Schematics of *α*_5_*β*_1_ integrin in its bent-closed, inactive state with FN fragment 7-10 unbounded (left), extended-active state in complex with FN (middle), and under an applied load (right). B) The Cryo-EM structure of *α*_5_*β*_1_-FN with the individual integrin heads and FN fragments labeled. The MD simulations applied a velocity to the P1142 residue while restraining K559 and E36. Zoomed in region shows wildtype synergy site with R1374 and R1379 (left) and double mutated R1374/9A synergy site (right). D154 binds to R1379 and is shown as a reference. SYN: synergy site. RGD: arg-gly-asp.

Wildtype and mutated structures were solvated in a TIP3P water box (18nm x 45nm x 19nm) with 0.15mM NaCl. Energy was minimized for 15k steps with the steepest gradient descent algorithm, followed by an equilibration sequence of a 1ns NVT simulation at 310K followed by a 10ns NPT simulation at 1 bar and 310K, per physiological conditions. Equilibration was verified by ensuring that the RMSD of the fully unrestrained complexes (Figure S1) were within 0.3nm resolution of cryoEM.

The K559 and E36 residues at the proximal ends of the integrin headpieces were then restrained. P1142 at the distal end of the FN fragment was pulled at 10 and 1nm/ns using a 50kJ/mol/nm spring with an umbrella potential for 3ns and 20ns, respectively. The steered MD simulations used a 2fs timestep. We visualized the crystal structures and MD simulation trajectories using Visual Molecular Dynamics (VMD) 1.9.4a (29). All parameters for the MD simulations are available in the supplementary materials (Table S1). The force and extension at *α*_5_*β*_1_-FN’s center-of-mass (COM) were derived directly from the output files from Gromacs. The extension was measured as the displacement of the *α*_5_*β*_1_-FN’s center-of-mass with respect to the first simulation frame. The radius of gyration of the *α*_5_ and *β*_1_ heads was measured using the built-in Gromacs function, gmx gyrate. Distances between key bonds at R1374 and R1379 were calculated by averaging the distance between atom pairs that could form hydrogen bonds using the VMD bond select and graph tool. We used a distance cutoff of 0.35nm (3.5 Angstrom) and donor-hydrogen-acceptor angle cutoff of 30 in VMD to detect hydrogen bonds.

### Synergy Site Departure Energy

To calculate the energy required for the synergy site to depart from *α*_5_, we used in-house Python code to integrate the force and COM extension data from the beginning of the simulation to the time of the force peak just before the rapid increase in extension rate. Since the force-extension data is non-monotonic, we first fitted a piece-wise linear function over the force-extension data before integrating with trapezoid rule.

### Force Distribution Analysis

Time-resolved force distribution analysis (trFDA) was used to measure the punctual stresses based on the Coulombic interactions at all residues across all simulation time steps (30). The punctual stress is the absolute value of scalar pairwise forces exerted on each residue. Normally, stress would be in units of energy. However, the developers of punctual stress defined it as “force on a dimensionless point” which uses units of force (kJ/mol-nm). We opted to use this definition of punctual stress to remain consistent with past studies. Parameters for the trFDA are available in the supplementary materials (Table S2).

### Long-term NPT Equilibration Simulations

Longer term stability of the *α*_5_*β*_1_-FN complex after synergy site mutagenesis was tested with two 250ns NPT simulations of *α*_5_*β*_1_-FN9-10: one wildtype and one R1374/9A mutant. The 7NWL pdb file was truncated from *α*_5_*β*_1_-FN7-10 to *α*_5_*β*_1_-FN9-10. A R1374/9A double mutation was again induced in silico via MODELLER 10.4 (28). The system contained ≈ 1.3M atoms in a 15nm x 30nm x 30nm box after solvation. NaCl concentration was kept at 0.15mM. The 250ns NPT simulation was preceded by a 15k step energy minimization and 1ns NVT as described previously. 100kJ/mol-nm^2^ restraints were placed on residues D603, E445, and D1328 (Figure 4A) in the x and y directions, representing the remaining structures of integrin and FN while limiting periodic box crossing. No other restraints were placed. We used GROMACS 2020.4 (26) to measure backbone RMSDs, nonbonded energies, axes of inertia, distances, and hydrogen bonds. Axes of inertia were used to calculate angles by taking the inverse cosine of the dot product of a unit vector pair. Measurements were tested for normality with the Kolmogorov-Smirnov test. Since all data was non-normal, the wildtype and mutant trajectories were compared using the Wilcoxon signed rank test (*α* = 0.05).

### Extensional Stiffness of *α*_5_ and *β*_1_ headpieces

Extensional stiffnesses of *α*_5_ and *β*_1_ headpieces were determined independently using 100ns NPT simulations. The 7NWL pdb file was isolated to either the *α*_5_ head (≈ 438K atoms in a 16.5nm x 16.5nm x 16.5nm box post solvation) or *β*_1_ head (≈ 463K atoms in a 16.8nm x 16.8nm x 16.8nm box post solvation). Again, energy minimization for 15k steps and a 1ns NVT as previously described were run in GROMACS prior to the 100ns NPT simulation. Extensional stiffness, *k*, for each molecule was calculated using:

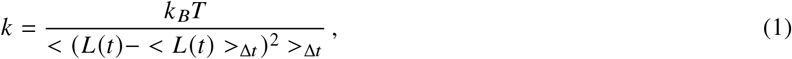

where *k*_*B*_*T* is Boltzmann’s constant, *T* = 310*K, L* (*t*) is the length of the reaction coordinate at time, *t*, and <> denotes the time average (31). For *α*_5_, the center-of-mass distance between D154 (synergy site binding residue) and D603 (connects to lower integrin legs) in *α*_5_ was chosen as the length of the reaction coordinate. Similarly for *β*_1_, the Metal-Ion Dependent Adhesion Site (MIDAS; binds to RGD) and E445 (connects to lower integrin legs) were chosen. After the system had equilibrated, we used the latter 50ns for the extensional stiffness calculation. For each molecule, the distance data was divided into five 10ns blocks. Distances were saved every 10ps during the simulation, resulting in 1000 data points per block to calculate five *k* values per head. A Wilcoxon signed-rank test compared the means of the extensional stiffnesses of *α*_5_ and *β*_1_. The angle between the propeller and thigh in *α*_5_ was measured as described previously.

### Whole-cell Finite Element Model

We used a whole-cell FE model to calculate the *α*_5_*β*_1_-FN concentration and force in a wildtype and mutant cell. We have previously modeled the cell-substrate interface using a whole-cell FE model; we refer the reader to that publication for the full set of model equations (23). In the present work, we introduced key changes to the catch bond model. We modeled the cell as a 2D elastic disk with neo-Hookean constitutive material properties on a rigid substrate,

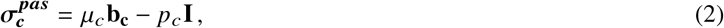

where 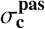 is the passive cell stress. The cell shear modulus is, *μ*_*c*_=1kPa (32, 33). The deformation was characterized by the left Cauchy-Green tensor **b**_**c**_. The pressure *p*_*c*_ was computed from plane stress boundary conditions.

An isotropic active stress field was applied inside the cell to model cell contractility,

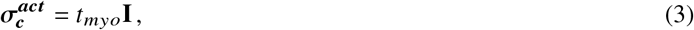

where 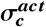 is the active cell stress due to an actomyosin traction, *t*_*myo*_ in Pa (33, 34):

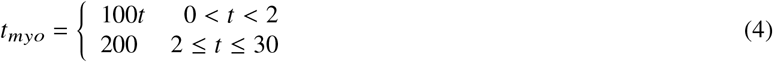

where *t* is the simulation time.

We used an existing catch bond model of adhesion to calculate the force-dependent concentration of *α*_5_*β*_1_-FN bonds per node in the FE mesh (35–38). The catch model assumed that the *α*_5_*β*_1_-FN complexes behave as parallel springs that connect and disconnect to the substrate based on an association constant, *K*_*on*_ and on a force dependent dissociation constant, *K*_*off*_, respectively.

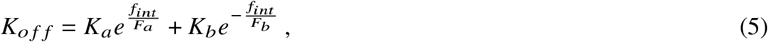

where *K*_*a*_, *F*_*a*_, *K*_*b*_, and *F*_*b*_ are fitted parameters (Table S3) adapted from Bidone et al (38) and Takagi et al (39). *f*_*int*_ is the magnitude of the force per *α*_5_*β*_1_-FN bond. The force vector per bond, (**f**_**int**_), is computed via the *α*_5_*β*_1_-FN spring constant *k*_*int*_ = 0.5pN/nm (17) and the spring extension vector **u**_**int**_:

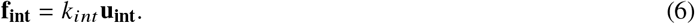

The force per node, **f**_*i,node*_ is related to the dimensionless concentration of *α*_5_*β*_1_-FN bonds *C* with respect to the maximum bond density *ρ*_*i,max*_ = 100*μ*m^2^ (40), and the local adhesion area *A* at that node,

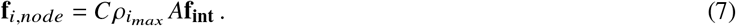

At any node, *i* given the previous value of the bond concentration, *C*, the updated bond concentration *C*_*t*+Δ*t*_ at each progressive time step is

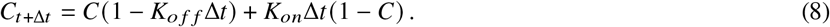

Note that the updated eq. (8) is based on treating the bond kinetics in the limit of an ordinary differential equation discretized in time with an explicit Euler scheme.

The internal force balance for the cell includes the elastic cell deformation 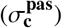 and the active cell contractile stress 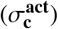:

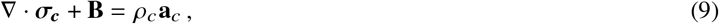

in which 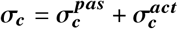 is the total cell stress, **B** is the total body force on the cell, *ρ*_*c*_ = 1000kg/m^3^ is the cell density (41) and **a**_*c*_ is the cell acceleration.

The strong form of the elastodynamic equation 9 has boundary conditions of the form **σ**·**n** = **t** on boundary Γ_*c*_, which includes the external forces on the circumference. Assuming 2D plane stress, the body forces on the cell arise from *α*_5_*β*_1_-FN bond forces and viscous drag forces. The internal forces were computed through the weak form. Briefly, we multiplied equation 9 by test function, *v*, integrated over a domain Ω_*c*_ of thickness 1*μ*m, and applied divergence theorem to get the following weak form for the cell.

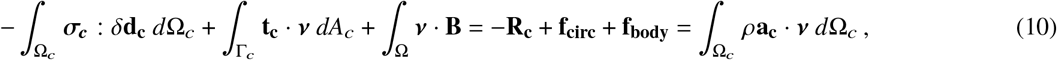

The δ**d**_**c**_ is the variation of the symmetric velocity gradient, i.e. virtual work by moving each node by an independent variation *v*. **R**_**c**_ is the residual (internal forces) and the external force acting at a node of the cell mesh is composed of the forces on the circumference, **f**_**circ**_ and the forces on the body, **f**_**body**_:

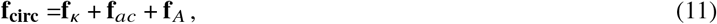

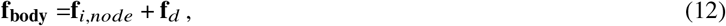

where **f**_*i,node*_ is the force due to *α*_5_*β*_1_-FN at each node, **f**_*d*_ is viscous drag, **f**_*k*_ is curvature regularization, **f**_*ac*_ is a random fluctuation at the cell boundary from actin polymerization, and **f**_*A*_ is an area penalty to counteract cell contractility.

The mesh was updated by a dynamic explicit mesh generator, El Topo (42), during the simulation run. The explicit mid-point rule was used for time integration of the second order system of equations to update nodal velocities and positions. The whole-cell FE simulation ran with a time step of 50*μ*s over the course of an assigned time of *t*_*sim*_ = 30s. There were a total of three simulation runs per R1374/9A mutant and wildtype catch bond condition, respectively. The three simulation bond concentration and force outputs were time-averaged per condition.

## RESULTS AND DISCUSSION

### FN9-*α*_5_ disengagement coincides with synergy site deactivation

We analyzed force-extension in conjunction with punctual stress to determine the role of the synergy site in FN9-*α*_5_ disengagement. The initial force-extension curve of the wildtype *α*_5_*β*_1_-FN structure followed a linear response for both 10 and 1 nm/ns pull rates until peaking at 729pN and 462pN, respectively (Figure 2A and B). The peak forces coincided with sharp decreases in the punctual stress at the synergy site, namely at sites R1374 and R1379 in FN9. R1379 has been shown to be connected to D154 in the *α*_5_ head via a salt bridge (13). However, R1374 has not been previously observed to be actively linked to *α*_5_. At both pull rates, R1374 retained higher punctual stresses than R1379, but the sequence of disengagement was dependent on the pull rate. Under the faster pull rate condition, the salt bridge was disrupted prior to a reduction in force on *α*_5_*β*_1_-FN and punctual stress at R1374. This indicated that while the load on FN was sufficient to overcome the energetic barrier to break the salt bridge connecting FN to *α*_5_, persistent electrostatic interaction at R1374 enabled FN9 to remain near the *α*_5_ head. This was not observed under the slower pull rate simulation, where we noted simultaneous punctual stress reduction in R1374 and R1379 at the peak force time point. While the punctual stresses at both residues were elevated during load ramping, synergy site engagement reduced after the force peak.

**Figure 2.**
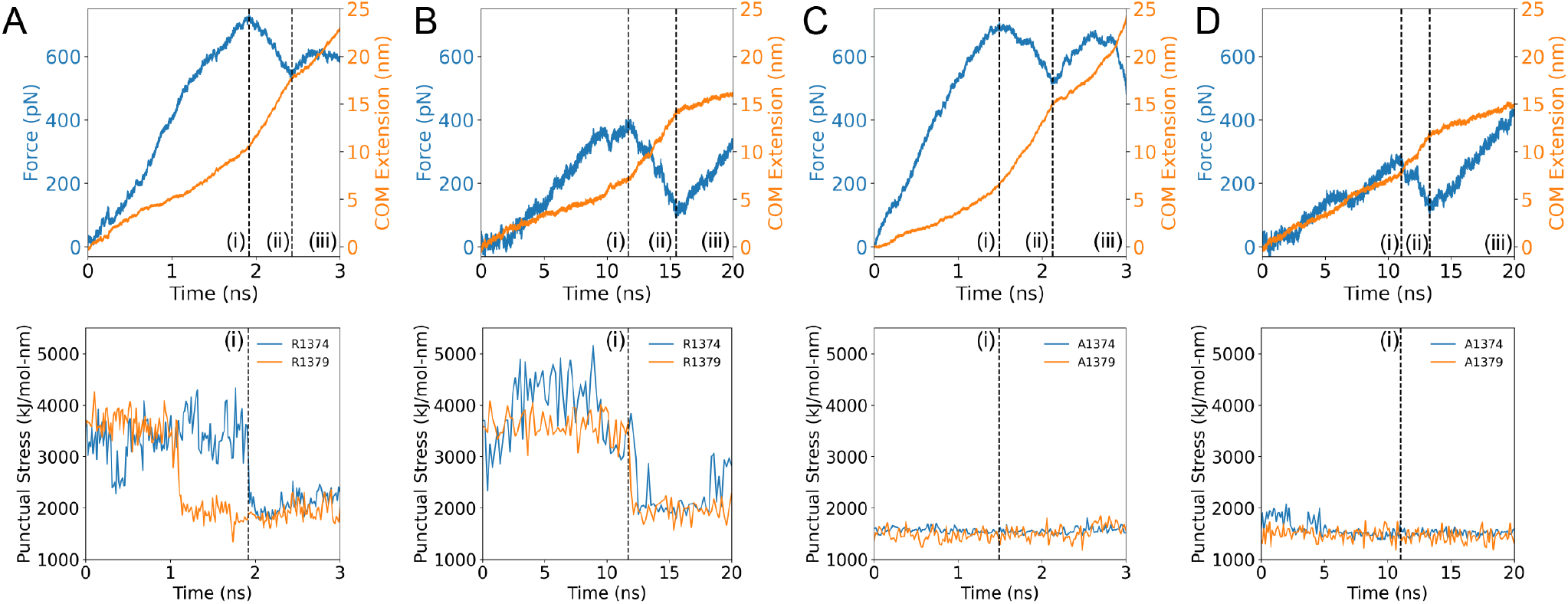
Force and COM extension over time plotted over punctual stress at R1374/1379 of the synergy site for A) 10nm/ns wildtype *α*_5_*β*_1_-FN, B) 1nm/ns wildtype *α*_5_*β*_1_-FN, C) 10nm/ns R1374/9A *α*_5_*β*_1_-FN, and D) 1nm/ns R1374/9A *α*_5_*β*_1_-FN. Positions (i), (ii), and (iii) correspond to the time at the peak force, local minimum, and final frame, respectively.

R1374 and R1379 were contributors to punctual stress at the synergy site prior to the drop in force on *α*_5_*β*_1_-FN (Figure S2). In both pull rate conditions, the combined punctual stress at R1374/9 prior to the force peak was on average two times higher than other synergy site residues. Due to the high electrostatic activity of both sites prior to FN9 and *α*_5_ separation, we mutated both residues (R1374/9A) to evaluate their roles in maintaining *α*_5_*β*_1_-FN’s structural response to force. At 10nm/ns, the force response of the wildtype and mutant *α*_5_*β*_1_-FN were similar, peaking at 729pN and 704pN, respectively (Figure 2C).

However, the punctual stresses at A1374 and A1379 were 45% and 40% lower in the mutant case than the wildtype (Figure 2C and D), indicating that the mutation disrupted synergy site engagement, but not necessarily reduced force transmission. Similar trends were observed in the 1nm/ns force rate condition, where the punctual stresses at A1374 and A1349 were small relative to R1374 and R1379, and the first peak force was lower in the mutant case (wildtype = 462pN, mutant = 291pN; Figure 2D).

Although our results appeared to conflict with the understanding that synergy site mutagenesis decreases cell adhesion strength, the relative energetic barrier required to separate the synergy site from integrin revealed closer agreement with the literature (17, 18, 24, 39, 43). While we noticed a 171pN difference (37% less than the wildtype) in the first peak force in the 1nm/ns mutant model, we only noted a 25pN drop (3% less than the wildtype) in the 10nm/ns model. This is likely a consequence of the high pull rates used in these models that may hide molecular mechanisms. Therefore, long term simulations at slower pull rates and smaller forces are needed to overcome this limiting factor. We worked towards this goal in a later section. For now, to overcome this potential conflict with the literature, we opted to use the area under the force-extension curve (Figure S4) as a proxy for measuring synergy site departure energy, which would be related to the energy barrier required to pull FN9 away from *α*_5_. We defined the *synergy site departure force* as point (i) in all simulations (Figure 2). Forces recorded after the *synergy site departure force* would work to unfold FN and unbind RGD. We found that the synergy site departure energies were greater in the wildtype, in line with past in vitro experiments that show greater binding affinity of *α*_5_*β*_1_ integrin to FN in the presence of the synergy site (24, 39). At 10nm/ns, the wildtype and mutant energies were 4012pN-nm and 2715pN-nm, respectively. At 1nm/ns, the wildype and mutant had a energies of 1529pN-nm and 883pN-nm, respectively. These values do not have any physical meaning, but enabled a comparison between the wildtype and mutant. From our current steered MD data, we cannot make claims about the effect of the synergy site on RGD binding specifically. Free energy methods such as FEP (free energy perturbation) and MMPBSA (Molecular Mechanics Poisson-Boltzmann Surface Area) would be more appropriate to study these effects computationally and are the subject of ongoing work.

Punctual stress measurements provided insight into per-residue interactions at the synergy site and are substantiated by atomic-level interactions. Specifically, the formation and breakage of hydrogen bonds between *α*_5_ and FN9 are essential for relaying force between the two. Since high punctual stresses were observed on R1374 and R1379, we tracked bonds between R1379—D154 and R1374—E124 (Figure S5A). At both pull rates, the R1379—D154 salt bridge was broken before the maximum force was reached, while residue R1374 remained bounded to either E124 or E81 depending on the pull rate (Figure S5B-C). The measured distance between R1374—E124 was within the range of a hydrogen bond (0.35nm) after the departure of the R1379—D154 bond (10nm/ns case; Figure S5D). At the slower pull rate, R1374 transitioned from E124 to E81, maintaining contact between FN9 and *α*_5_*β*_1_ together with R1379—D154 (Figure S5E). Both bonds then released and the force on *α*_5_*β*_1_-FN consequently dropped. The R1374/9A double mutation severed the main points of contact between FN9 and *α*_5_*β*_1_, pushing the distance between the residues to 0.65nm, beyond the 0.35nm hydrogen bond length cutoff (Figure S5F).

For all test cases, the peak forces were followed by sharp increases in extension rate, suggesting a rapid conformational change of *α*_5_*β*_1_-FN (Figure 2). In the case of the wildtype 10nm/ns pull rate, the measured extension rate increased from 5.10nm/ns to 14.4nm/ns. Similarly, the wildtype 1nm/ns pull rate increased in rate from 0.547nm/ns to 1.82nm/ns (Table S4). Notably, there was a mismatch between the input rate and measured rate. Steered MD simulations attempt to control the pull rate via a virtual spring connecting a dummy atom to the pulled site. While the atom moves at a constant rate, the molecule’s response depends on the virtual spring deflection and local conformational changes associated with the molecule. Therefore, it is unlikely that the input pull rate matches the measured pull rate experienced by the molecule. Further, the output extension was measured as the distance traveled by *α*_5_*β*_1_-FN’s COM, which depends on the structural behavior.

Our reported forces and pull rates are many orders of magnitude higher than what has been tested using atomic force microscopy (AFM; 1 - 15 *μ*m/s) (43). Given our large 1.5M atom system, we compromised on the simulation time scale by applying extension rates within the bounds of past steered MD simulations of integrin (0.1 - 10 nm/ns) (22, 44). The fast extension rates contributed to simulated forces beyond what has previously been measured experimentally (single molecule rupture forces of 80-120pN) (43). Nevertheless, the difference between the forces generated at 1 and 10 nm/ns hinted at force-dependent behavior arising from synergy site engagement. Larger conformational changes were visually noted in the *α*_5_ head during 10nm/ns pulling compared to 1nm/ns pulling. Further, the mutants showed little to no changes in the movement of the *α*_5_ head, suggesting that the interactions at the synergy site could work to deform *α*_5_. Therefore, we quantified the conformational changes associated with synergy site engagement when subjected to high pull rates.

### Conformational response of *α*_5_ and *β*_1_ was hampered by lack of synergy site engagement

We informed the differences in force and extension rates across conditions by visualizing the structural changes of *α*_5_*β*_1_-FN under both pull rates for the wildtype and mutant cases. We used the radius of gyration to quantify conformational changes within *α*_5_ and *β*_1_ heads, with smaller radii indicating more compact proteins. In both wildtype runs, the *α*_5_ head, which is connected to the synergy site on FN9, stretched further than the *β*_1_ head, which is connected to the RGD motif on FN10. However, pull rate affected the degree of *α*_5_ stretching. The lower 1nm/ns pull rate resulted in 0.165nm increase in *α*_5_’s radius of gyration (Figure 3A), compared to a 0.407nm increase in the 10nm/ns rate simulation (Figure S3A). Most of the *α*_5_ head deformation resulted before the peak force and synergy site disengagement. For the respective 10nm/ns and 1nm/ns rates, 97.7% and 99.0% of the max *α*_5_ head deformation occurred prior to the peak force, when the synergy site loosened. From the observations of *α*_5_*β*_1_-FN’s quaternary structure, we noticed the *α*_5_ head straightening while FN9 remained connected at the synergy site (Figure 3C). Further, at higher forces, *α*_5_ underwent a greater degree of stretching while FN9 unfolded (Figure S3C). In contrast, lower forces seemed to encourage synergy site disengagement prior to FN unfolding. Our observation suggests that *α*_5_*β*_1_-FN’s catch bond dynamics may be promoted by greater synergy site interaction in combination with *α*_5_ extension to resist larger forces. The greater interaction may stem from the hydrogen bond electrostatics at R1374 and R1379 that bridge *α*_5_ to FN9 (Figure S5).

**Figure 3.**
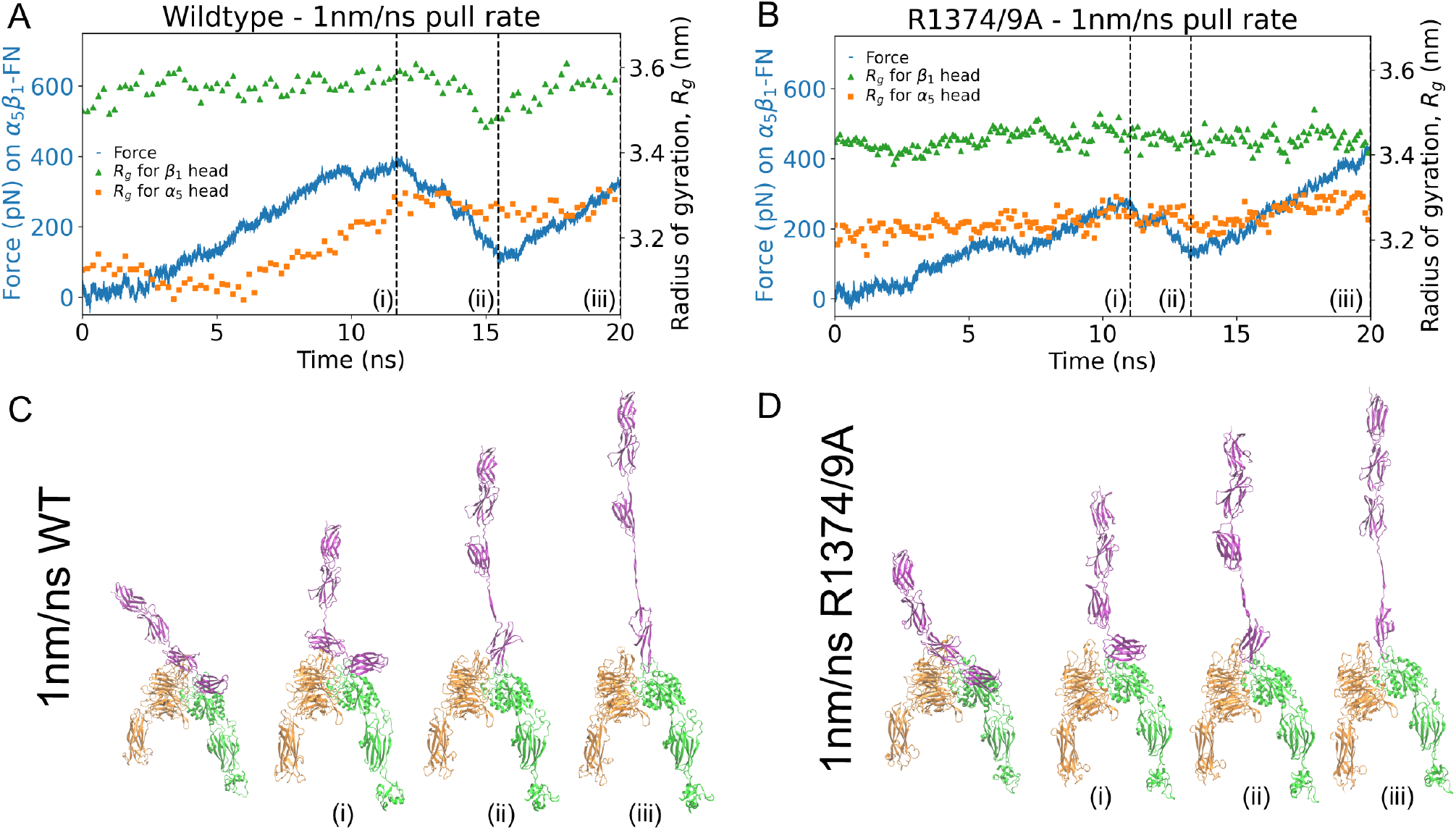
Force on *α*_5_*β*_1_-FN and radius of gyration of *α*_5_ and *β*_1_ head for the 1nm/ns runs for the A) wildtype and B) mutant. Positions (i), (ii), and (iii) correspond to the time at the peak force, local minimum, and final frame. The four shown frames from the simulation correspond to the first frame, (i) peak force, (ii) local minimum, and (iii) final frame for C) wildtype and D) mutant.

**Figure 4.**
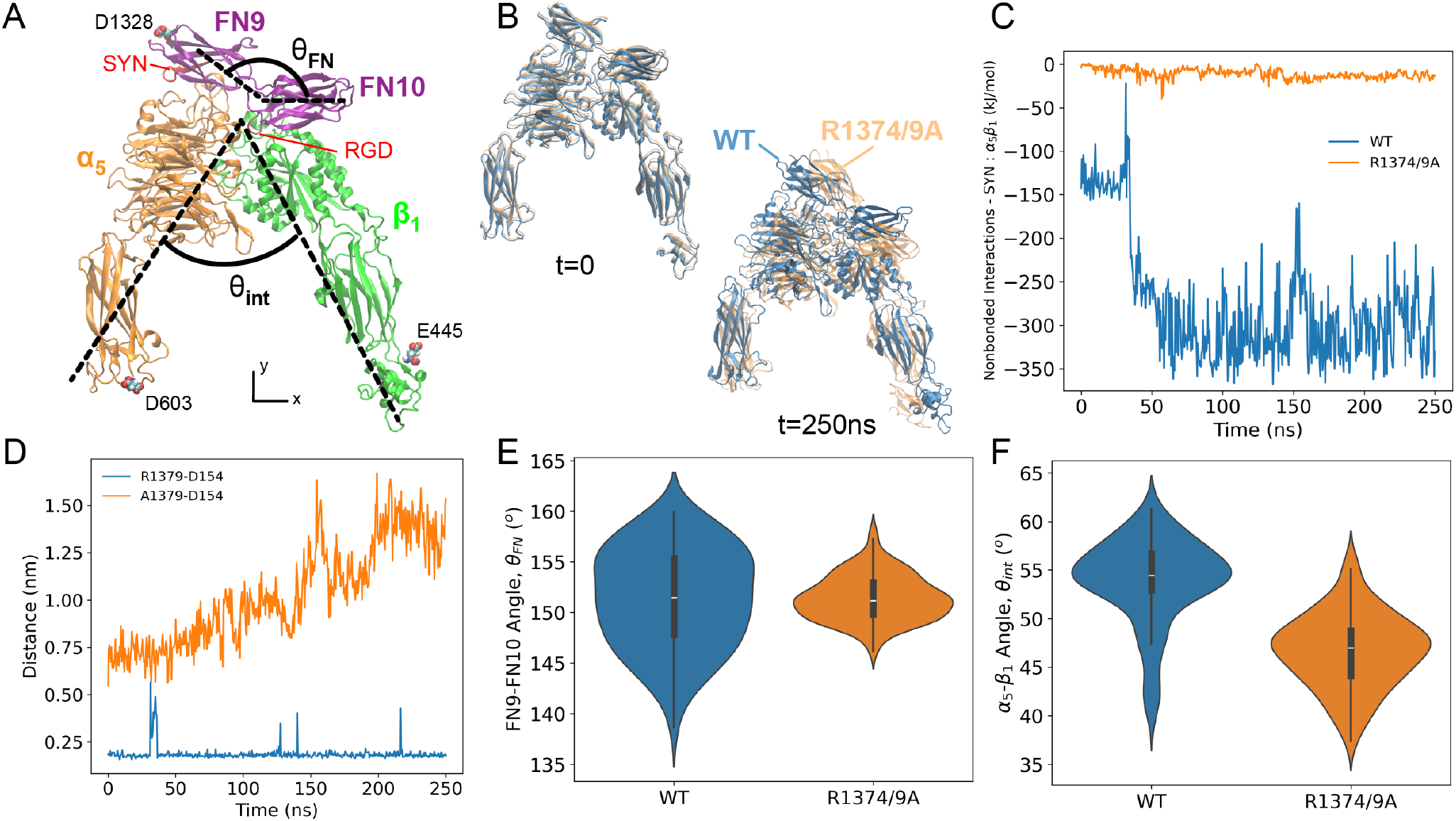
A) Cryo-EM structure *α*_5_*β*_1_-FN9-10. Small restraints were placed on D603, E445, and D1348 in the x and y directions to mimic the respective continuing structures of integrin and FN. *θ*_*int*_ was defined the angle between the principal axes of inertia of *α*_5_ and *β*_1_, respectively. Similarly, *θ*_*FN*_ was defined as the angle between the principal axes of inertia of FN9 and FN10, respectively. Dashed lines are hand-drawn and indicate an approximation of the principal axes. SYN: synergy site. B) Superposition of the wildtype (blue) and mutant (orange) during the first and last frames of the respective 250ns simulations. C) Nonbounded interaction energy between the synergy site and *α*_5_*β*_1_ integrin for wildtype and mutant. D) Minimum distance between residue 1379 (FN9) and D154 (*α*_5_) for wildtype and mutant. E) Violin plot of FN9-10 angle for last 50ns of 250ns simulation (WT = 151.4 ± 4.9°, R1374/9A = 151.4 ± 2.2°, *p* = 0.98). F) Violin plot of *α*_5_*β*_1_ angle for last 50ns of 250ns simulation (WT = 53.9 ± 4.3°, R1374/9A = 46.7 ± 3.8°, *p* < 0.0001).

We tested the degree to which the synergy site contributed to structural changes in *α*_5_*β*_1_-FN by mutating the site (R1374/9A) and again measuring the radius of gyration of *α*_5_ and *β*_1_ under an external load on FN. Surprisingly, the mutant pulled at 10nm/ns still resulted in conformational changes of the *α*_5_ head, with the radius of gyration increasing by 0.266nm. However, this was less than the 0.407nm increase observed in the wildtype (Figure S3B). Further, the mutant pulled at the slower 1nm/ns showed virtually no deformation of *α*_5_ or *β*_1_ (Figure 3B). Investigating the quaternary structure of the mutant revealed that FN9 was separated immediately from *α*_5_ (Figures 3D and S3D). As the FN beta sheets stacked vertically in alignment with the pulling direction, the force increased and peaked as soon as FN10 begun to unfold. For all simulations, the *β*_1_ head kept a more stable conformation, maintaining its radius of gyration within 0.12nm. These results are indicative of a new mechanism whereby *α*_5_ and FN deformation patterns may be altered due to interactions at the synergy site. However, the fast pull rates are five orders of magnitude higher than even the slowest AFM pull rates, posing the question of whether these states may be realized and more importantly, have a physical meaning. So, while our results were promising, we aimed to address the pull rate limitation by conducting longer term simulations and emphasizing our analysis on the synergy site and integrin interaction.

### Synergy site interactions maintained FN9 and *α*_5_ close

We used two 250ns NPT simulations of *α*_5_*β*_1_ integrin in complex with FN9-10 (wildtype and R1374/9A) to understand the role of the synergy site in maintaining integrin and FN conformational stability. Visual observation showed separation of mutant FN9 away from integrin as well as minor deviations to the integrin headpieces (Figure 4B). Therefore, we investigated the connection between FN9 and integrin. As expected, we found that the nonbonded interactions (van der waals and coulombic energies) between the synergy site and *α*_5_*β*_1_ were greater in the wildtype structure (Figure 4C). These results aligned with the shorter distance between R1379 in FN9 and D154 in *α*_5_ (Figure 4D) as well as the greater number of hydrogen bonds between the synergy site and *α*_5_ (Figure S6A).

Lower synergy site engagement widened the gap between FN9 and *α*_5_, but only minor structural changes in the integrin heads and FN were realized. We conducted structural analyses using the final 50ns of the 250ns simulation. The nonbonded interactions (Figure 4A), the hydrogen bond count (Figure S6A), and backbone RMSD (Figure S6B) of *α*_5_*β*_1_-FN9-10 (wildtype and mutant) leveled off at ≈ 200ns, suggesting system equilibration. Longer simulations would be necessary to evaluate whether the system fully equilibrated, but based on these initial trends, we enforced the latter 50ns cutoff. Since the synergy site in FN9 and RGD in FN10 are two anchoring contact points for integrin, we posited that releasing FN9 from *α*_5_ via synergy site inhibition would increase FN9-10 flexibility. Interestingly, the means of the FN9-10 angles (*θ*_*FN*_) in both cases was not statistically significant and variance was greater in the wildtype (Figure 4E), which would indicate that the wildtype FN9-10 was fluctuating to a greater degree even as the synergy site was interacting more strongly. Further, the *α*_5_-*β*_1_ angle (*θ*_*int*_) in the wildtype was 7.2° larger than the mutant, pointing to a modest closing of the integrin heads in the mutant (Figure 4F). This closing was predominantly a result of FN9-10 rotation rather than a state transition of *α*_5_. The propeller-thigh angle (*θ* _*α*5_) was 4.7° greater in the mutant, whereas the *β*_1_-FN10 angle (*θ*_*β*1−*FN*10_) was 12.1° lower in the mutant (Figure S8). FN9-10 retained its shape, with only a 0.01nm difference in radius of gyration between mutant and wildtype (Figure S7A-B). Additionally, there was no statistically significant difference in the radius of gyration of *α*_5_ between mutant and wildtype (Figure S7C-D). The radius of gyration of *β*_1_ in the mutant was 0.16nm smaller (Figure S7E-F), indicating a small amount of compression of *β*_1_ as it interacted with FN10. The time series data of *θ*_*FN*_ (Figure S6C), *θ*_*int*_ (Figure S6D), *θ* _*α*5_ (Figure S8B), and *θ*_*β*1−*FN*10_ (Figure S8D) showed overlap between mutant and wildtype throughout the entire simulation, meaning that some states may be similar to each other, but on average, the conformational measurements suggest that the synergy site locks FN9 to *α*_5_ and prevents rotation of FN9-10.

The unlocking of FN9 due to reduced synergy site energetics did not promote appreciable changes at integrin’s RGD binding location. We first measured the nonbonded interaction energies between RGD and *α*_5_*β*_1_, including the MIDAS cation, which showed no differences in energies after, and even before the imposed 200ns cutoff (Figure S9A). Additionally, the number of hydrogen bonds between *α*_5_ and RGD (Figure S9B) well as *β*_1_ and RGD (Figure S9C) were similar between the wildtype and mutant. From this data, we assumed that RGD would be a stable location for FN to maintain binding to integrin regardless of synergy site engagement. To confirm the conformational stability at the RGD binding area, we measured the mean and minimum distances between notable interactions at this site (Figure S10A). These included RGD-MIDAS (Figure S10B-C), D227-RGD (Figure S10D-E), and S134-MIDAS (Figure S10F-G). As expected, the distances between these pairs remained small in both the wildtype and mutant. Although there were differences in the S134-MIDAS mean and minimum distance, the observed 0.05-0.75nm distance difference was not enough to decrease the absolute interaction energy at the mutant’s RGD site (Figure S9A). The stability of the RGD binding site enabled it to behave like a pivot point for mutated FN9-10 when FN9 dislodged from the synergy site. Since our data suggests that RGD remained stable regardless of synergy site engagement, we reasoned that the additional synergy site interaction energies in the wildtype would only bolster *α*_5_*β*_1_-FN binding. From past in vitro experiments, RGD alone is known to be sufficient to support some *α*_5_*β*_1_ integrin binding and cell adhesion, though it has been shown that the synergy site promotes longer lasting binding and stronger cell adhesion when it binds in tandem with RGD to secure FN (6, 24). The synergy site alone does not support cell adhesion as well as only RGD, or both RGD and the synergy site (45, 46), which may be attributed to the synergy site’s lower nonbonded interaction energy (Figure 4C) compared to RGD (Figure S9A). However, as mentioned, free energy methods must be considered to include the entropic effects that we do not account for in this work.

Collectively, our observations of the 250ns NPT trajectories support the conjecture that the synergy site reinforces integrin engagement with the matrix (13, 24). Further, our accelerated steered MD models imply that force between the synergy site and *α*_5_ integrin head may induce conformational changes of *α*_5_ integrin. Overall, our results highlight the importance of the synergy site clip in stabilizing and reinforcing the *α*_5_*β*_1_-FN bond after initial catch bond formation, which has also been previously suggested experimentally (9, 25, 47, 48). While cell adhesion can be negated altogether by an RGD deletion as demonstrated by spinning disk assays, the R1374/9A double mutation reduces cell adhesion strength by around 90% (24). So, while adhesion could still occur, the bond strength was compromised due to the synergy site mutation, which has also been shown previously through single molecule AFM (43). Additionally, past surface plasmon resonance binding assays measure an 11-fold decrease in affinity between *α*_5_*β*_1_ and R1374A FN compared to wildtype (39). Clearly, the role of the synergy site in maintaining a firm adhesion cannot be understated. Here, we propose how the synergy site may give rise to specific molecular states of *α*_5_*β*_1_-FN, since it holds FN9 near *α*_5_. Our steered MD models at a 1nm/ns pull rate showed a decrease in initial synergy site departure energy after mutagenesis, implying that there is a greater energetic barrier in breaking the synergy site than when it is inhibited. Further, the 1nm/ns wildtype model predicts that the connection between FN9-*α*_5_ maintained by the synergy site could deform the *α*_5_ head when loaded, which was not observed in the 1nm/ns mutant run. While our MD study highlighted the reinforcing role of the synergy site at the molecular scale, we also sought to explore how this adhesion reinforcement may dynamically manifest at the whole cell scale.

### Synergy site presence led to adhesion reinforcement by recruiting *α*_5_*β*_1_ integrin

We employed a whole-cell FE model that analyzed the adhesion interface that contained *α*_5_*β*_1_-FN bonds under an isotropic cell contraction that drove bond extension (Figure 5A). Our simple model demonstrated an adaptive reinforcement of collective *α*_5_*β*_1_-FN bonds due to the stronger binding affinity afforded by the synergy site. We modified the parameters for the *α*_5_*β*_1_-FN binding kinetics (Table S3) to produce bond lifetime curves for the wildtype bond and R1374/9A mutant (Figure 5B). The differences in parameters between the two bond types resulted in an 11-fold decrease in *α*_5_*β*_1_-FN bond concentration (Figure 5C), but no increase in equilibrium force (Figure 5D). The areas of high concentrations and high forces are present at the periphery of the cell model during contraction (Figures 5E and 5F), which has been shown by 2D Fluorescence Resonance Energy Transfer (FRET) and traction force microscopy (TFM) assays (25). Notably, mutant bonds compensate for the lack of number of bonds by sustaining more of the cell’s contractile load. The higher recruitment of wildtype bonds distributes the forces more evenly across the cell model.

**Figure 5.**
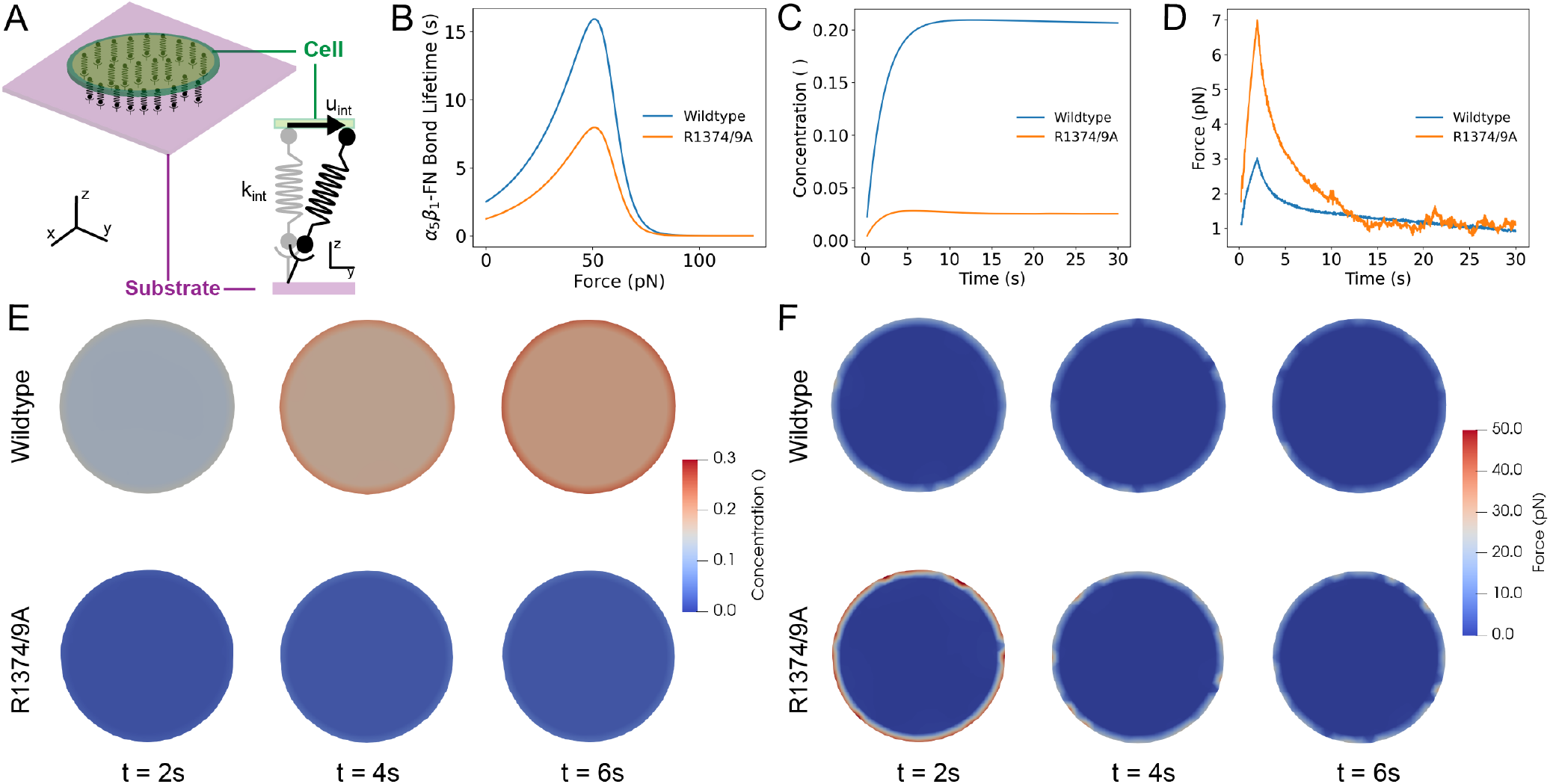
A) Schematic of whole-cell interface model that assumes that integrin behaves as a spring that is stretched due to cell contraction. B) Catch bond model: *α*_5_*β*_1_-FN bond lifetime versus applied force for wildtype (adapted from (38, 39)). C) Concentration over time of wildtype and mutant *α*_5_*β*_1_-FN. D) Force over time of wildtype and mutant *α*_5_*β*_1_-FN. E) Frames at times 2, 4, and 6s indicating the concentration of *α*_5_*β*_1_-FN bonds across the cell-substrate interface during a 200Pa uniform contraction. F) Frames at times 2, 4, and 6s indicating the distribution of *α*_5_*β*_1_-FN bond force across the cell-substrate interface during a 200Pa uniform contraction.

Our whole-cell FE model sheds light on the dynamic force balance at short timescales that are not as apparent experimentally. TFM of cells plated on 2D substrates have shown that cell contraction and individual bond force were not altered due to an absence of the synergy site (25, 48). Our model used the same 200Pa cell contraction across both conditions, but showed a stark difference in how the adhesion forces are handled by the bonds. Namely, while forces eventually equalized between mutant and wildtype conditions, we observed an initial dynamic adjustment of high forces at the cell model’s boundary for mutant bonds (Figure 5F). Specifically, average forces measured from mutated bonds peaked at 7pN, while wildtype bonds peaked at 3pN; both average bond forces were within the previously measured 1-7pN range (25). A body of work has shown the reduction in cell adhesion strength at the single molecule and whole cell scale due to a lack of synergy site engagement (24, 25, 43, 48). In spite of the reduced bond strength, our model showed that, under minimal tension, the binding affinity gain due to the presence of the synergy led to a more stable, dynamic force balance across the *α*_5_*β*_1_-FN bonds on the cell model’s surface.

### Pivot-clip mechanism of *α*_5_*β*_1_-FN as a model for cell adhesion reinforcement

The mechanosensitive pivot-clip mechanism provides a model to consider how the *α*_5_*β*_1_-FN catch bond reinforces cell adhesion across molecular and cell scales under cell-matrix forces (Figure 6). Long term NPT simulations indicated that role of the synergy site was to clip FN9 close to *α*_5_ as evidenced by the increased separation between FN9 and *α*_5_ in the mutant. The dislodging of FN9 did not modify the stability of the RGD site. In our steered MD simulations, for both pull rates tested in the wildtype *α*_5_*β*_1_-FN, the unbinding of FN9-*α*_5_ coincided with a plateauing of *α*_5_ extension (Figures 3A and S3A). With the link between FN9-*α*_5_ broken, FN10 was free to rotate about the RGD motif on *β*_1_ (Figures 3D and S3D). The FN10 rotation about the RGD site was maintained in the mutant steered MD runs while diminishing the increase in radius of gyration of *α*_5_ (Figure S3B and D). Based on the structural changes observed on *α*_5_ in the steered MD simulations, the synergy site clipped the *α*_5_ head to FN9 while the RGD motif on *β*_1_ acted as a pivot for FN10 (Figure 6). Since *α*_5_ preferentially stretched instead of *β*_1_, we conducted 100ns NPT simulations of each integrin head to measure each of their relative extensional stiffness. Upon confirming a stable RMSD after 50ns (Figure S11A), we averaged the measured *α*_5_ and *β*_1_ head distances over five 10ns blocks (Figure S11B) to quantify extensional stiffness. We measured extensional stiffnesses of 2587 pN/*μ*m and 174548 pN/*μ*m for the *α*_5_ and *β*_1_ heads, respectively (Figure S11C). Based on the distance fluctuations, *β*_1_ remained more static, while *α*_5_ seemed to oscillate. We also found that the propeller-thigh angle of *α*_5_ decreased (Figure S11D), giving *α*_5_ a more bent shape (Figure S11E). We reasoned that the link between the propeller and thigh grants *α*_5_ its flexibility to stretch when force is applied, while *β*_1_’s rigidity could provide a route for forces to transmit towards cytoskeletal proteins. While it has been known that the synergy site plays a role in catch bond dynamics (17, 24), the clip engagement under force could be one mechanism by which the synergy site enables catch bond dynamics at the molecular scale. Using our pivot-clip model (Figure 6), forces generated at the cell-matrix interface would need to first overcome the synergy site clip energy barrier. In parallel, *α*_5_ would resist forces by stretching prior to FN9 unclipping, also leading to a higher barrier than if the synergy site were not present. Additionally, the rigidity of *β*_1_ could facilitate downstream mechanosignaling via talin. Namely, talin binds to the *β*_1_ tail and has been shown to be a mechanosensitive protein that interacts with vinculin and focal adhesion kinase to promote focal adhesion maturation and nuclear localization of transcriptional coregulator, Yes-Associated Protein (36, 49, 50). However, larger forces could also increase the probability of FN unbinding from *α*_5_*β*_1_, especially when the additional energetic barrier from the synergy site is not present. Past assays have demonstrated that *α*_5_*β*_1_-FN unbinding occurs with greater likelihood when the synergy site is inhibited; moreover, *α*_5_*β*_1_-FN losing its catch bond characteristics (18, 24). To determine the exact pathway of the force transmission across the *α*_5_*β*_1_-FN catch bond with and without the synergy site, much longer and slower MD simulations are needed. Along those lines, more investigation is warranted to elucidate how the full structure of *α*_5_*β*_1_ dynamically couples with mechanosensitive cytoskeletal proteins at the atomistic scale.

**Figure 6.**
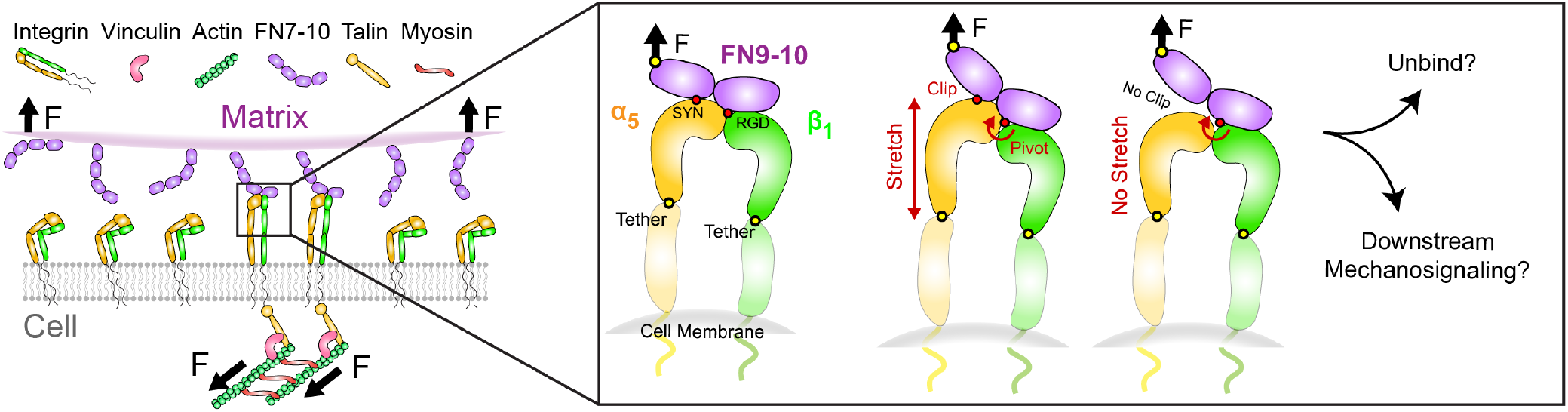
Proposed model for synergy site clip engagement leading to *α*_5_ deformation during mechanosensing while RGD acts as a pivot for FN10. In this model, force transmits across the clip, stretching *α*_5_. The additional energetic barrier provided by the clip could afford *α*_5_*β*_1_-FN greater resistance to unbinding. The rigidity of *β*_1_ relative to *α*_5_ may allow for force transmission across the membrane and towards the mechanosensitive cytoskeletal protein, talin, leading to downstream mechanosignaling.

In the context of outside-in signaling, the *α*_5_*β*_1_-FN pivot-clip mechanism demonstrates how the synergy site could route force via *β*_1_ towards mechanosignaling proteins in the cytoplasm, like talin, leading to integrin clustering. According to the outside-in activation model, integrins maintain a bent-closed, low affinity state before undergoing a conformational change to an extended, active conformation upon encountering an ECM ligand (Figure 1A) (51–53). In contrast, the inside-out model proposes that the adaptor protein talin would bind to the cytoplasmic tail of integrin, allowing it to activate and subsequently bind to its ligand (51–53). While the current hypothesis states that binding between FN and *α*_5_*β*_1_ triggers an opening of integrin’s cytoplasmic tails leading to an accumulation of adaptor proteins that resist cell-matrix forces (Figure 6), further studies are needed to elucidate the mechanism behind integrin activation. Multiple steered MD models have been employed to interrogate *β*_3_ integrin activation (22, 44, 54–57), with few investigating the cytoplasmic end of *β*_1_ integrin (58, 59). However, to our knowledge, our approach is unique in that we model the interface between FN and the *α*_5_*β*_1_ integrin heads, where forces are transmitted bidrectionally between the cell and its matrix.

Our study acknowledges several limitations. Firstly, we made the assumption that the proximal ends of the integrin heads were anchored by fully extended integrin legs tightly held by tails in the cell membrane. While this assumption contributed to model stability, it is worth noting that the head-leg junction has been suggested to possess greater flexibility (13). Relaxing the constraints on the proximal ends to allow lateral movement may introduce flexibility without the added complexity of integrating the legs. Secondly, our steered MD models applied a large, vertical pulling rate. While this approach is advantageous for directly stressing the points of contact between FN and *α*_5_*β*_1_, it could introduce biased pulling and rotational forces that are unrealistic, which would decrease model confidence. Multiple runs and a parametric study of boundary conditions must be considered when confirming our MD simulations in future works investigating tension or other loading modalities, such as shear or torsion. Lastly, our focus was on a specific integrin subtype. The intricate nature of cell-matrix interactions involves multiple integrin subtypes and their respective ligands. Due to the prohibitive cost of molecular dynamics simulations, alternative approaches such as coarse-grained or agent-based models, capable of examining cell-matrix interactions at a broader systems level and over extended timescales, may be necessary.

## CONCLUSION

This work advances our understanding of cell mechanobiology by introducing a mechanosensitive mechanism, termed pivot-clip, by which *α*_5_*β*_1_ integrin reinforces cell adhesion. Using FE and MD simulations, we shed light on a biophysical connection between the cell and ECM that underpins many cellular behaviors that drive physiology and pathology. Critically, we also demonstrated binding domains that promote catch bond dynamics in the context of cell-matrix mechanosensing. Looking forward, we envision elucidating how the force-dependent, pivot-clip mechanism interacts with its surrounding machinery and how it may be transformed via novel therapeutics. As our understanding of cell adhesion progresses, we aim to develop informed approaches to target diseases that rely on transmitting forces via cell-matrix bonds.

## AUTHOR CONTRIBUTIONS

A.R.M., A.B.T., and M.R.K.M. conceptualized and designed the research. A.R.M., A.B., and W.W. performed the research and analyzed data. A.R.M. wrote the manuscript. G.D.O., A.B.T., and M.R.K.M. supervised the research and edited the manuscript.

### ACKNOWLEDGMENTS

This research used Expanse at San Diego Supercomputer Center through allocation MCB100146 from Advanced Cyberinfrastructure Coordination Ecosystem: Services & Support (ACCESS) super-computing facilities, supported by the National Science Foundation (NSF) (Grant Nos. 2138259, 2138286, 2138307, 2137603, and 2138296). We also acknowledge the additional allocation, BIO230214 for use of Expanse as well as the Negishi Community Cluster managed by the Rosen Center for Advanced Computing. A.R.M. was supported by the Ford Predoctoral Fellowship, UC Berkeley College of Engineering Robert N. Noyce Fellowship, and Hearts to Humanity Eternal Research Fellowship. A.B. was supported by the National Institute of Health Maximizing Access to Research Careers T34 Award. W.W. was supported by the NSF Research Experience for Undergraduates via the Transfer-to-Excellence Program at University of California, Berkeley. A.B.T was supported by the UC Berkeley Miller Institute for Basic Research in Science. We thank members of the Molecular Cell Biomechanics and Berkeley Biomechanics Labs for fruitful discussions that led to the improvement of this manuscript. Special thank you to Mohammad Khavani for his suggestions to improve the molecular dynamics simulations.

## REFERENCES

1. Damiano, J. S., A. E. Cress, L. A. Hazlehurst, A. A. Shtil, and W. S. Dalton, 1999. Cell adhesion mediated drug resistance (CAM-DR): role of integrins and resistance to apoptosis in human myeloma cell lines. Blood, the Journal of the American Society of Hematology 93:1658–1667.

2. Bachmann, M., S. Kukkurainen, V. P. Hytönen, and B. Wehrle-Haller, 2019. Cell adhesion by integrins. Physiological reviews 99:1655–1699.

3. Lee, M. H., P. Ducheyne, L. Lynch, D. Boettiger, and R. J. Composto, 2006. Effect of biomaterial surface properties on fibronectin–α5β1 integrin interaction and cellular attachment. Biomaterials 27:1907–1916.

4. Hynes, R. O., 2002. Integrins: bidirectional, allosteric signaling machines. cell 110:673–687.

5. Ross, T. D., B. G. Coon, S. Yun, N. Baeyens, K. Tanaka, M. Ouyang, and M. A. Schwartz, 2013. Integrins in mechanotransduction. Current opinion in cell biology 25:613–618.

6. Cutler, S. M., and A. J. García, 2003. Engineering cell adhesive surfaces that direct integrin α5β1 binding using a recombinant fragment of fibronectin. Biomaterials 24:1759–1770.

7. Scheiblin, D. A., J. Gao, J. L. Caplan, V. N. Simirskii, K. J. Czymmek, R. T. Mathias, and M. K. Duncan, 2014. Beta-1 integrin is important for the structural maintenance and homeostasis of differentiating fiber cells. The international journal of biochemistry & cell biology 50:132–145.

8. Humphrey, J. D., E. R. Dufresne, and M. A. Schwartz, 2014. Mechanotransduction and extracellular matrix homeostasis. Nature reviews Molecular cell biology 15:802–812.

9. Benito-Jardón, M., S. Klapproth, I. Gimeno-LLuch, T. Petzold, M. Bharadwaj, D. J. Müller, G. Zuchtriegel, C. A. Reichel, and M. Costell, 2017. The fibronectin synergy site re-enforces cell adhesion and mediates a crosstalk between integrin classes. Elife 6:e22264.

10. Hou, J., D. Yan, Y. Liu, P. Huang, and H. Cui, 2020. The roles of integrin α5β1 in human cancer. OncoTargets and therapy 13329–13344.

11. Zhao, X.-K., Y. Cheng, M. Liang Cheng, L. Yu, M. Mu, H. Li, Y. Liu, B. Zhang, Y. Yao, H. Guo, et al., 2016. Focal adhesion kinase regulates fibroblast migration via integrin beta-1 and plays a central role in fibrosis. Scientific reports 6:19276.

12. Cao, L., J. Nicosia, J. Larouche, Y. Zhang, H. Bachman, A. C. Brown, L. Holmgren, and T. H. Barker, 2017. Detection of an integrin-binding mechanoswitch within fibronectin during tissue formation and fibrosis. ACS nano 11:7110–7117.

13. Schumacher, S., D. Dedden, R. V. Nunez, K. Matoba, J. Takagi, C. Biertümpfel, and N. Mizuno, 2021. Structural insights into integrin α5β1 opening by fibronectin ligand. Science Advances 7:eabe9716.

14. Kechagia, J. Z., J. Ivaska, and P. Roca-Cusachs, 2019. Integrins as biomechanical sensors of the microenvironment. Nature Reviews Molecular Cell Biology 20:457–473.

15. Schaffner, F., A. M. Ray, and M. Dontenwill, 2013. Integrin α5β1, the fibronectin receptor, as a pertinent therapeutic target in solid tumors. Cancers 5:27–47.

16. Cox, D., M. Brennan, and N. Moran, 2010. Integrins as therapeutic targets: lessons and opportunities. Nature reviews Drug discovery 9:804–820.

17. Kong, F., A. J. García, A. P. Mould, M. J. Humphries, and C. Zhu, 2009. Demonstration of catch bonds between an integrin and its ligand. Journal of Cell Biology 185:1275–1284.

18. Strohmeyer, N., M. Bharadwaj, M. Costell, R. Fässler, and D. J. Müller, 2017. Fibronectin-bound α5β1 integrins sense load and signal to reinforce adhesion in less than a second. Nature Materials 16:1262–1270.

19. Dansuk, K. C., and S. Keten, 2021. Self-strengthening biphasic nanoparticle assemblies with intrinsic catch bonds. Nature communications 12:85.

20. Dansuk, K. C., S. Pal, and S. Keten, 2023. A catch bond mechanism with looped adhesive tethers for self-strengthening materials. Communications Materials 4:60.

21. Yuan, Z., X. Duan, X. Su, Z. Tian, A. Jiang, Z. Wan, H. Wang, P. Wei, B. Zhao, X. Liu, et al., 2023. Catch bond-inspired hydrogels with repeatable and loading rate-sensitive specific adhesion. Bioactive Materials 21:566–575.

22. Kulke, M., and W. Langel, 2020. Molecular dynamics simulations to the bidirectional adhesion signaling pathway of integrin αVβ3. Proteins: Structure, Function, and Bioinformatics 88:679–688.

23. Montes, A. R., G. Gutierrez, A. Buganza Tepole, and M. R. K. Mofrad, 2023. Multiscale computational framework to investigate integrin mechanosensing and cell adhesion. Journal of Applied Physics 134:114702.

24. Friedland, J. C., M. H. Lee, and D. Boettiger, 2009. Mechanically activated integrin switch controls α5β1 function. Science 323:642–644.

25. Chang, A. C., A. H. Mekhdjian, M. Morimatsu, A. K. Denisin, B. L. Pruitt, and A. R. Dunn, 2016. Single molecule force measurements in living cells reveal a minimally tensioned integrin state. ACS nano 10:10745–10752.

26. Abraham, M. J., T. Murtola, R. Schulz, S. Páll, J. C. Smith, B. Hess, and E. Lindahl, 2015. GROMACS: High performance molecular simulations through multi-level parallelism from laptops to supercomputers. SoftwareX 1:19–25.

27. Schrödinger, L., 2015. The PyMOL Molecular Graphics System, Version 1.8.

28. Webb, B., and A. Sali, 2016. Comparative protein structure modeling using MODELLER. Current protocols in bioinformatics 54:5–6.

29. Humphrey, W., A. Dalke, and K. Schulten, 1996. VMD: visual molecular dynamics. Journal of molecular graphics 14:33–38.

30. Costescu, B. I., and F. Gräter, 2013. Time-resolved force distribution analysis. BMC biophysics 6:1–5.

31. Tong, D., N. Soley, R. Kolasangiani, M. A. Schwartz, and T. C. Bidone, 2023. Integrin αIIbβ3 intermediates: From molecular dynamics to adhesion assembly. Biophysical Journal 122:533–543.

32. Luo, Q., D. Kuang, B. Zhang, and G. Song, 2016. Cell stiffness determined by atomic force microscopy and its correlation with cell motility. Biochimica et Biophysica Acta (BBA)-General Subjects 1860:1953–1960.

33. Schierbaum, N., J. Rheinlaender, and T. E. Schäffer, 2019. Combined atomic force microscopy (AFM) and traction force microscopy (TFM) reveals a correlation between viscoelastic material properties and contractile prestress of living cells. Soft Matter 15:1721–1729.

34. Chen, C., J. Xie, L. Deng, and L. Yang, 2014. Substrate stiffness together with soluble factors affects chondrocyte mechanoresponses. ACS applied materials & interfaces 6:16106–16116.

35. Guo, Y., S. Calve, and A. B. Tepole, 2022. Multiscale mechanobiology: Coupling models of adhesion kinetics and nonlinear tissue mechanics. Biophysical Journal 121:525–539.

36. Cheng, B., W. Wan, G. Huang, Y. Li, G. M. Genin, M. R. Mofrad, T. J. Lu, F. Xu, and M. Lin, 2020. Nanoscale integrin cluster dynamics controls cellular mechanosensing via FAKY397 phosphorylation. Science advances 6:eaax1909.

37. Bell, G. I., 1978. Models for the specific adhesion of cells to cells: a theoretical framework for adhesion mediated by reversible bonds between cell surface molecules. Science 200:618–627.

38. Bidone, T. C., A. V. Skeeters, P. W. Oakes, and G. A. Voth, 2019. Multiscale model of integrin adhesion assembly. PLOS Computational Biology 15:e1007077.

39. Takagi, J., K. Strokovich, T. A. Springer, and T. Walz, 2003. Structure of integrin α5β1 in complex with fibronectin. The EMBO journal 22:4607–4615.

40. Gaudet, C., W. A. Marganski, S. Kim, C. T. Brown, V. Gunderia, M. Dembo, and J. Y. Wong, 2003. Influence of type I collagen surface density on fibroblast spreading, motility, and contractility. Biophysical journal 85:3329–3335.

41. Neurohr, G. E., and A. Amon, 2020. Relevance and regulation of cell density. Trends in cell biology 30:213–225.

42. Brochu, T., and R. Bridson, 2009. Robust topological operations for dynamic explicit surfaces. SIAM Journal on Scientific Computing 31:2472–2493.

43. Li, F., S. D. Redick, H. P. Erickson, and V. T. Moy, 2003. Force measurements of the α5β1 integrin–fibronectin interaction. Biophysical journal 84:1252–1262.

44. Chen, W., J. Lou, J. Hsin, K. Schulten, S. C. Harvey, and C. Zhu, 2011. Molecular dynamics simulations of forced unbending of integrin αVβ3. PLoS computational biology 7:e1001086.

45. Aota, S.-i., M. Nomizu, and K. M. Yamada, 1994. The short amino acid sequence Pro-His-Ser-Arg-Asn in human fibronectin enhances cell-adhesive function. Journal of Biological Chemistry 269:24756–24761.

46. Redick, S. D., D. L. Settles, G. Briscoe, and H. P. Erickson, 2000. Defining fibronectin’s cell adhesion synergy site by site-directed mutagenesis. The Journal of cell biology 149:521–527.

47. García, A. J., J. E. Schwarzbauer, and D. Boettiger, 2002. Distinct activation states of α5β1 integrin show differential binding to RGD and synergy domains of fibronectin. Biochemistry 41:9063–9069.

48. Tan, S. J., A. C. Chang, S. M. Anderson, C. M. Miller, L. S. Prahl, D. J. Odde, and A. R. Dunn, 2020. Regulation and dynamics of force transmission at individual cell-matrix adhesion bonds. Science Advances 6:eaax0317.

49. Zhou, D. W., M. A. Fernández-Yagüe, E. N. Holland, A. F. García, N. S. Castro, E. B. O’Neill, J. Eyckmans, C. S. Chen, J. Fu, D. D. Schlaepfer, et al., 2021. Force-FAK signaling coupling at individual focal adhesions coordinates mechanosensing and microtissue repair. Nature communications 12:2359.

50. Holland, E. N., M. A. Fernández-Yagüe, D. W. Zhou, E. B. O’Neill, A. U. Woodfolk, A. Mora-Boza, J. Fu, D. D. Schlaepfer, and A. J. García, 2024. FAK, vinculin, and talin control mechanosensitive YAP nuclear localization. Biomaterials 308:122542.

51. Jahed, Z., H. Shams, M. Mehrbod, and M. R. Mofrad, 2014. Mechanotransduction pathways linking the extracellular matrix to the nucleus. International review of cell and molecular biology 310:171–220.

52. Shattil, S. J., C. Kim, and M. H. Ginsberg, 2010. The final steps of integrin activation: the end game. Nature reviews Molecular cell biology 11:288–300.

53. Takagi, J., B. M. Petre, T. Walz, and T. A. Springer, 2002. Global conformational rearrangements in integrin extracellular domains in outside-in and inside-out signaling. Cell 110:599–611.

54. Driscoll, T. P., T. C. Bidone, S. J. Ahn, A. Yu, A. Groisman, G. A. Voth, and M. A. Schwartz, 2021. Integrin-based mechanosensing through conformational deformation. Biophysical journal 120:4349–4359.

55. Mehrbod, M., S. Trisno, and M. R. Mofrad, 2013. On the activation of integrin αIIbβ3: outside-in and inside-out pathways. Biophysical journal 105:1304–1315.

56. Su, S., Y. Ling, Y. Fang, and J. Wu, 2022. Force-enhanced biophysical connectivity of platelet β3 integrin signaling through Talin is predicted by steered molecular dynamics simulations. Scientific reports 12:4605.

57. Bidone, T. C., A. Polley, J. Jin, T. Driscoll, D. V. Iwamoto, D. A. Calderwood, M. A. Schwartz, and G. A. Voth, 2019. Coarse-grained simulation of full-length integrin activation. Biophysical journal 116:1000–1010.

58. Ji, Y., Y. Fang, and J. Wu, 2022. Tension Enhances the Binding Affinity of β1 Integrin by Clamping Talin Tightly: An Insight from Steered Molecular Dynamics Simulations. Journal of Chemical Information and Modeling 62:5688–5698.

59. Pan, D., and Y. Song, 2010. Role of altered sialylation of the I-like domain of β1 integrin in the binding of fibronectin to β1 integrin: thermodynamics and conformational analyses. Biophysical journal 99:208–217.

