## Supplementary Material for "Integrin Mechanosensing relies on Pivot-clip Mechanism to Reinforce Cell Adhesion"

Table S1: Steered Molecular Dynamics Parameters

| Parameter | Setting |
|---|---|
| Time step | 2fs |
| Number of steps (Time) for 1nm/ns | 10000000 (20ns) |
| Number of steps (Time) for 10nm/ns | 1500000 (3ns) |
| Integrator | Leapfrog algorithm |
| Constraint Algorithm | LINear Constraint Solver (LINCS) |
| Constraints | H-bonds constrained |
| Cutoff scheme | Verlet (Buffered neighbor searching) |
| Short-range neighbor list cutoff | 1.4nm |
| Short-range electrostatic cutoff | 1.4nm |
| Short-range van der Waals cutoff | 1.4nm |
| Electrostatics | Fast smooth Particle-Mesh Ewald (SPME) |
| Interpolation order | Cubic |
| Grid spacing for fast Fourier Transform | 0.12nm |
| Temperature coupling | Nosé-Hoover |
| Reference temperature | 310K |
| Temperature time constant | 1.0ps |
| Temperature coupled groups | Protein and non-protein |
| Pressure coupling | Off |
| Dispersion correction | long range dispersion corrections for energy and pressure |
| Velocity generation | Off |
| Harmonic potential | Umbrella |
| Force constant | 50 kJ/mol-nm$^2$ |
| Pull direction | y-direction (vertical) |
| Pull rate for 1nm/ns | 0.001nm/ps = 1nm/ns |
| Pull rate for 10nm/ns | 0.010nm/ps = 10nm/ns |

Table S2: Time resolved Force Distribution Analysis Parameter Settings

| Parameter | Setting |
|---|---|
| Pairwise forces | Summed |
| Pairwise groups | Protein |
| Residue based calculation | Punctual Stress |
| Pairwise force type | Coulombic interactions only |

Table S3: Catch bond parameters for whole-cell finite element model

| Variable | Wildtype | R1374/9A |
|---|---|---|
| $K_{on}$ | 0.1 $s^{-1}$ | 0.02 $s^{-1}$ |
| $K_a$ | 0.4 $s^{-1}$ | 0.8 $s^{-1}$ |
| $K_b$ | $4E-7s^{-1}$ | $8E-7s^{-1}$ |
| $F_a$ | $-25pN$ | |
| $F_b$ | $-15pN$ | |

Table S4: Measured extension rate (nm/ns) of $\alpha_5\beta_1$-FN under load in reference to the (i) peak and (ii) valley force as seen in Figure 2A-D

| | Prior to (i) | Between (i) and (ii) | After (ii) |
|---|---|---|---|
| 10nm/ns wildtype | 5.10 | 14.4 | 9.21 |
| 10nm/ns R1374/9A | 4.17 | 13.2 | 9.15 |
| 1nm/ns wildtype | 0.547 | 1.82 | 0.546 |
| 1nm/ns R1374/9A | 0.705 | 1.63 | 0.562 |

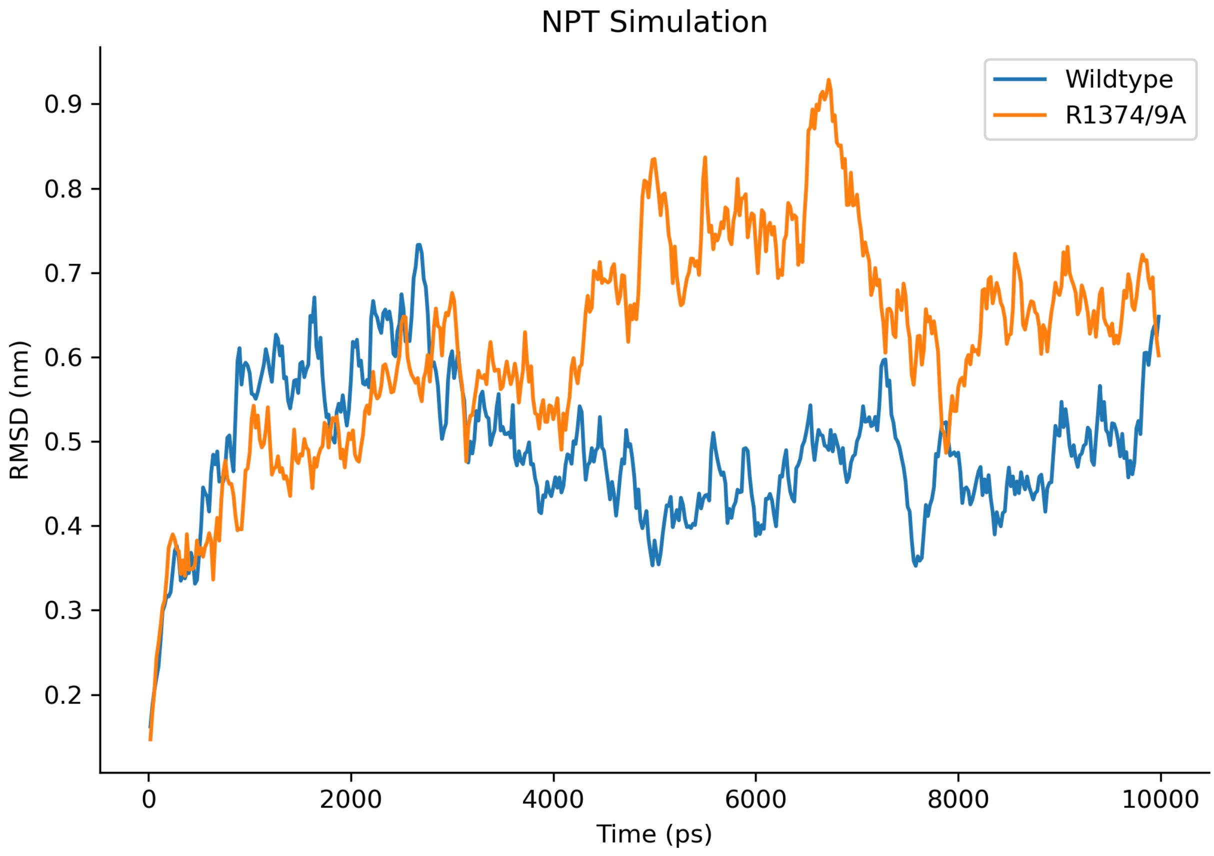


Figure S1: Root-mean-square deviation (RMSD) of wildtype and mutant during NPT simulation

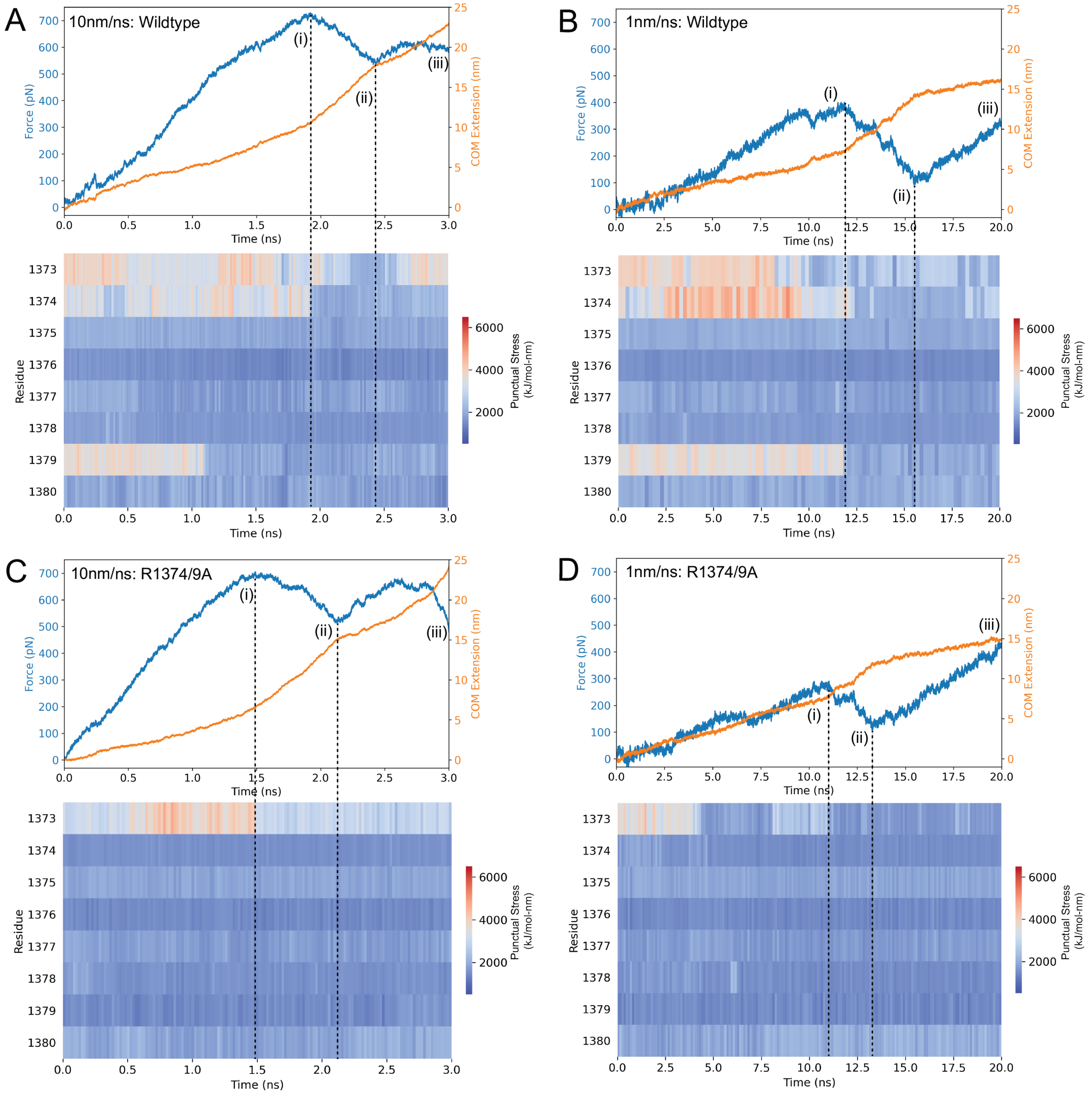


Figure S2: Force and COM extension over time plotted over punctual stress at the synergy site [1373-1380] for A) 10nm/ns wildtype $\alpha_5\beta_1$-FN, B) 1nm/ns wildtype $\alpha_5\beta_1$-FN, C) 10nm/ns R1374/9A $\alpha_5\beta_1$-FN, and D) 1nm/ns R1374/9A $\alpha_5\beta_1$-FN. Positions (i), (ii), and (iii) correspond to the time at the peak force, local minimum, and final frame, respectively.

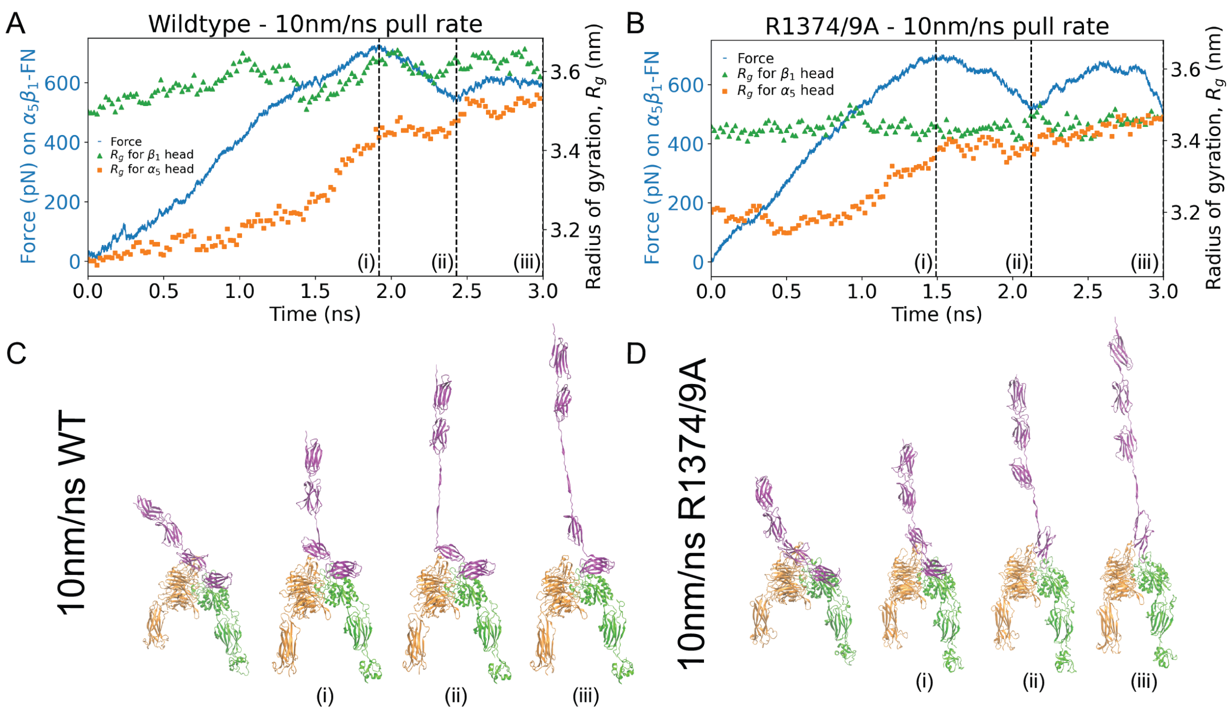


Figure S3: Force on $\alpha_5\beta_1$-FN and radius of gyration of $\alpha_5$ and $\beta_1$ head for the 10nm/ns runs for the A) wildtype and B) mutant. Positions (i), (ii), and (iii) correspond to the time at the peak force, local minimum, and final frame. The four shown frames from the simulation correspond to the first frame, (i) peak force, (ii) local minimum, and (iii) final frame for C) wildtype and D) mutant.

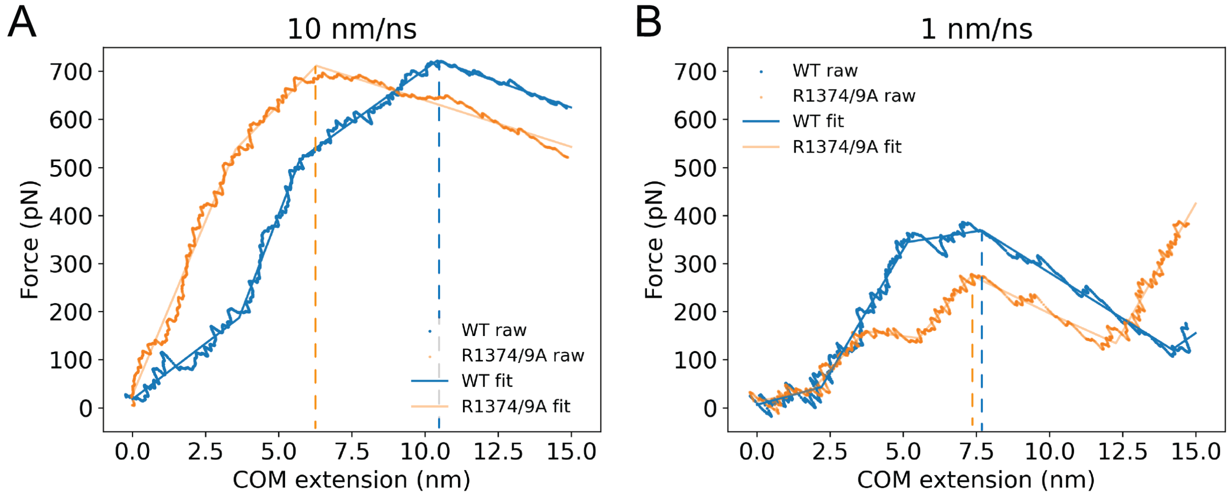


Figure S4: Force-extension plots for A) 10nm/ns and B) 1nm/ns pull rates. Dashed lines indicate the extension at the synergy site departure force. A moving average with a window size of 10ps was applied to the raw data for visualization purposes only. The piece-wise linear fit was performed over all points.

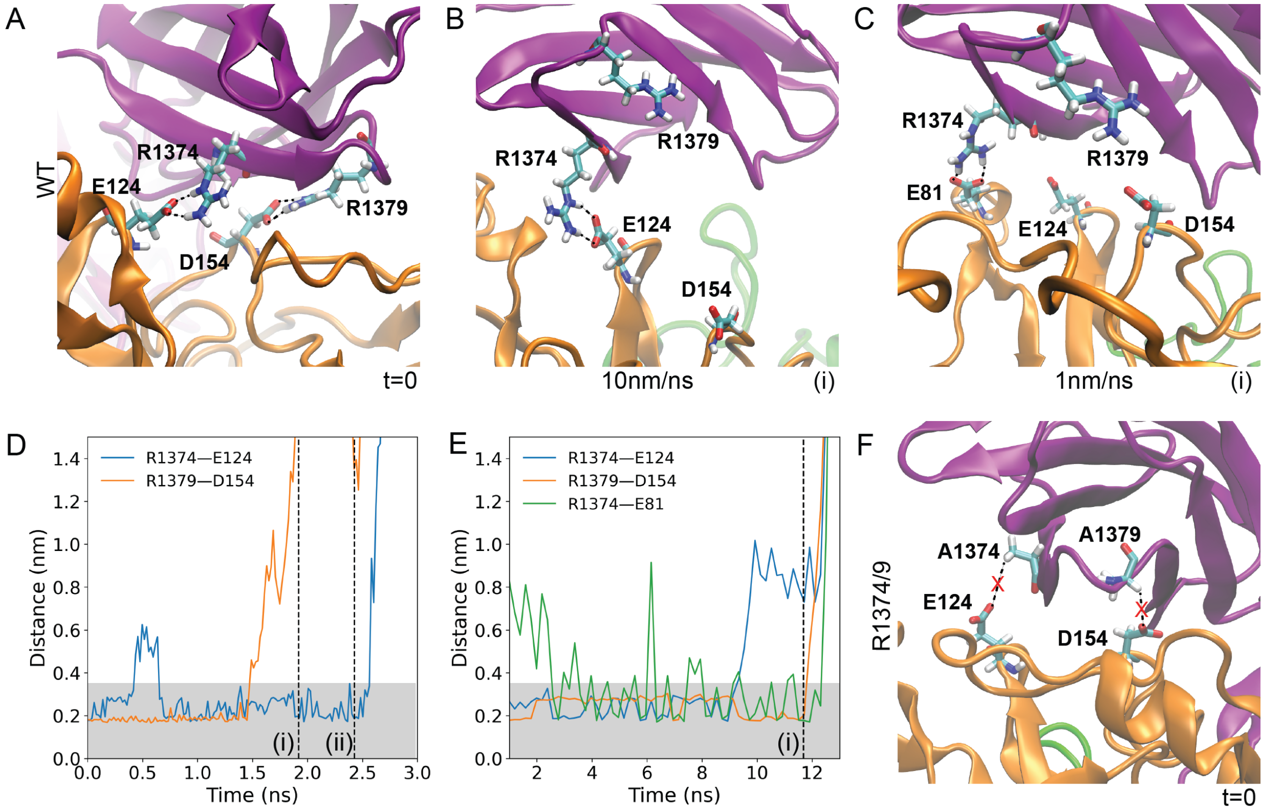


Figure S5: A) Interactions between R1374—E124 and R1379—D154 at the beginning of the wildtype simulations. B) At the force peak of the 10nm/ns wildtype simulation, the R1379—D154 salt bridge was broken but R1374—E124 remained. C) At the force peak of the 1nm/ns wildtype simulation, the R1379—D154 salt bridge was broken and R1374 formed a new hydrogen bond with E81. D) Distance between R1374—E124 and R1379—D154 for the 10nm/ns wildtype simulation. E) Distance between R1374—E124, R1379—D154, and R1374—E81. Shaded regions indicate 0.35nm, or the assumed approximate length for a hydrogen bond. The vertical dashed line is the time point of the force peak. F) The R1374/9A double mutation separated the A1374—E124 and A1379—D154 bonds to over 0.65nm, preventing hydrogen bond formation.

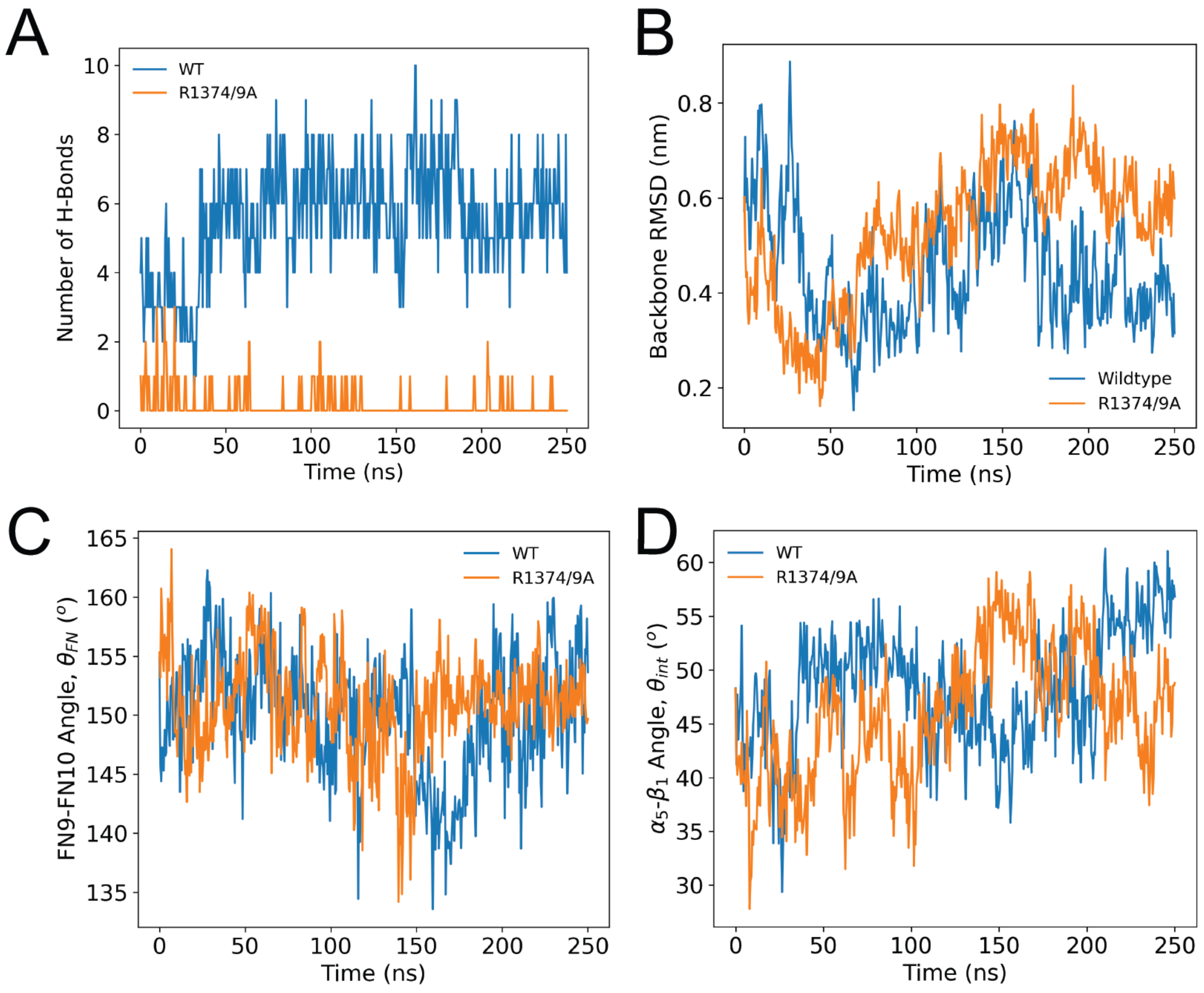


Figure S6: A) Number of hydrogen bonds, B) Backbone RMSD, C) FN9-FN10 Angle, and D) $\alpha$-$\beta_1$ angle over the 250ns simulation of the wildtype (WT) and mutant.

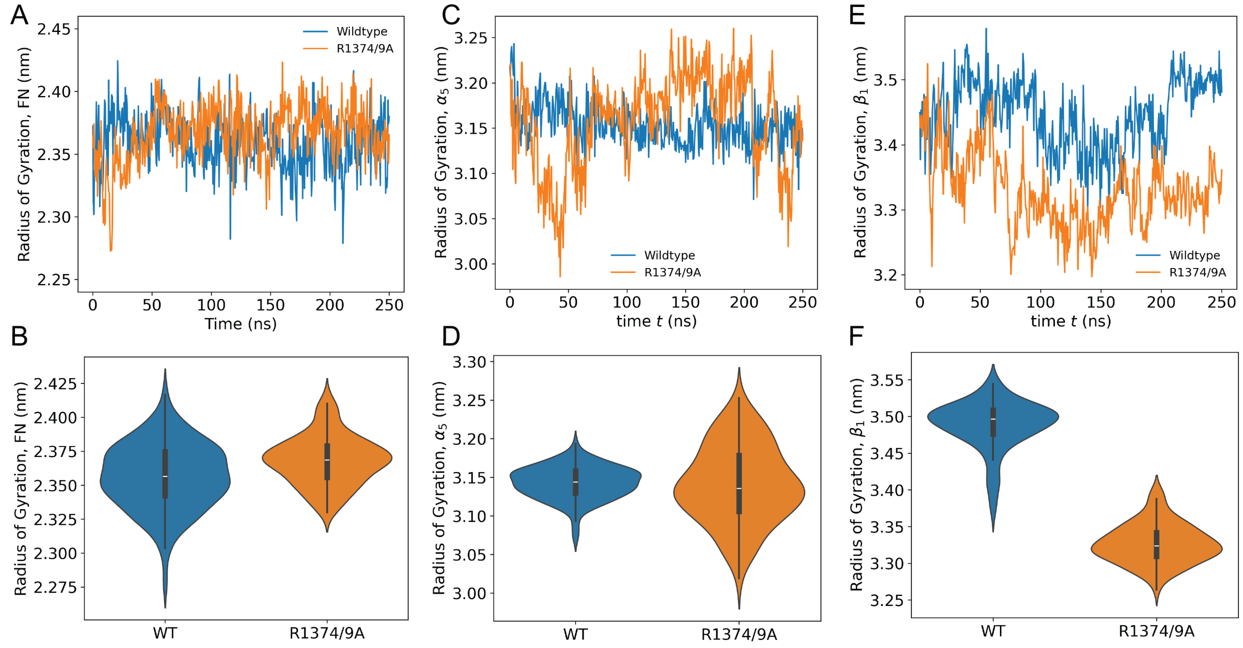


Figure S7: Time series data and violin plots of the radius of gyration of A-B) FN9-10 (WT $= 2.36 \pm 0.02$nm, R1374/9A $= 2.37 \pm 0.02$nm, $p = 0.0004$), C-D) the $\alpha_5$ head (WT $=3.14 \pm 0.02$nm, R1374/9A $= 3.14 \pm 0.05$nm, $p = 0.73$), and E-F) the $\beta_1$ head (WT $= 3.49 \pm 0.03$nm, R1374/9A $=3.33 \pm 0.03$pm, $p < 0.00001$)

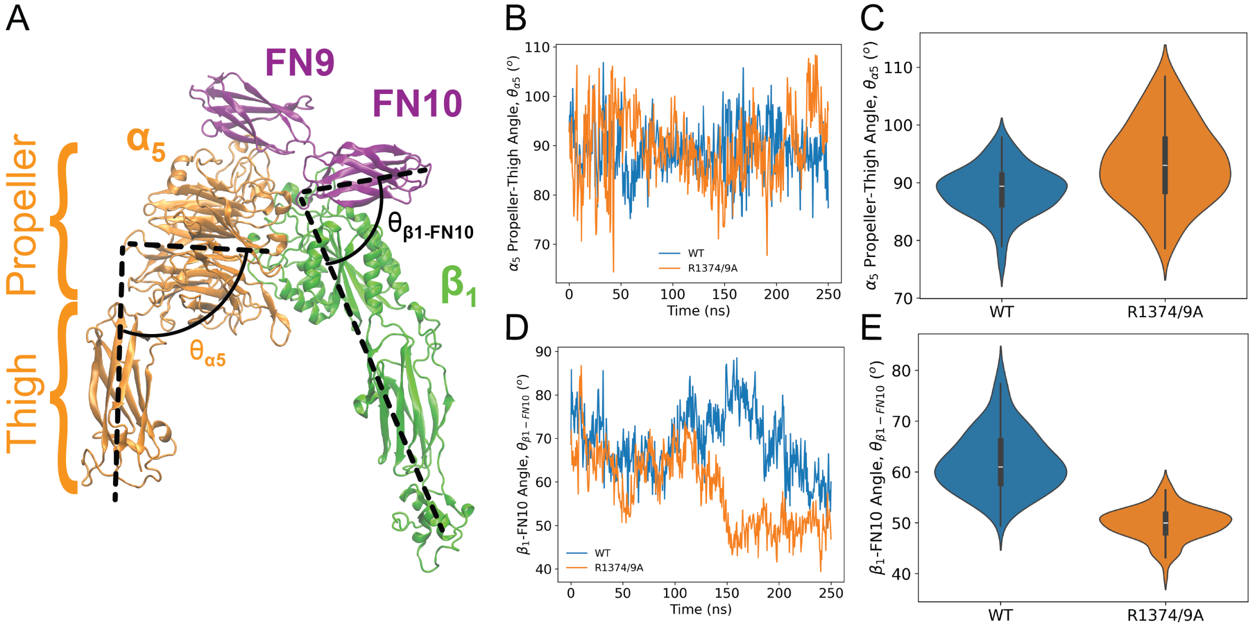


Figure S8: A) Cryo-EM structure of $\alpha_5\beta_1$-FN9-10 with the thigh propeller angle ($\theta_{\alpha5}$) and the $\beta_1$-FN10 angle ($\theta_{\beta1-FN10}$) labeled. B) Time series data of $\theta_{\alpha5}$. C) Violin plots of $\theta_{\alpha5}$ (WT $= 88.7 \pm 4.2°$, R1374/9A $= 93.4 \pm 6.8°$, $p = 0.3$). D) Time series data of $\theta_{\beta1-FN10}$. E) Violin plots of $\theta_{\beta1-FN10}$ (WT $= 62.0 \pm 6.0°$, R1374/9A $= 49.9 \pm 3.2°$, $p < 0.00001$).

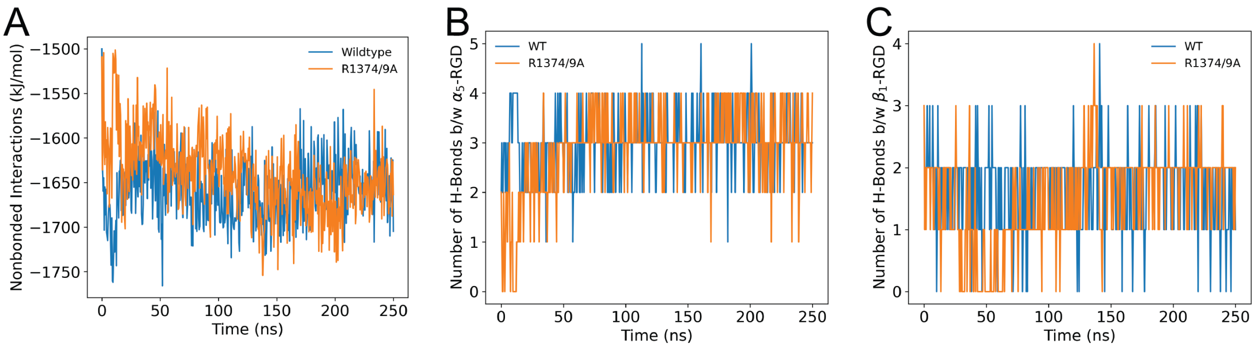


Figure S9: A) Nonbonded interactions at the RGD site which is the summation of the coulombic and van der waals energies for $\alpha_5$-MIDAS, $\alpha_5$-RGD, $\beta_1$-MIDAS, $\beta_1$-RGD, and RGD-MIDAS. B) Number of H-bonds between $\alpha_5$ and RGD. C) Number of H-bonds between $\beta_1$ and RGD.

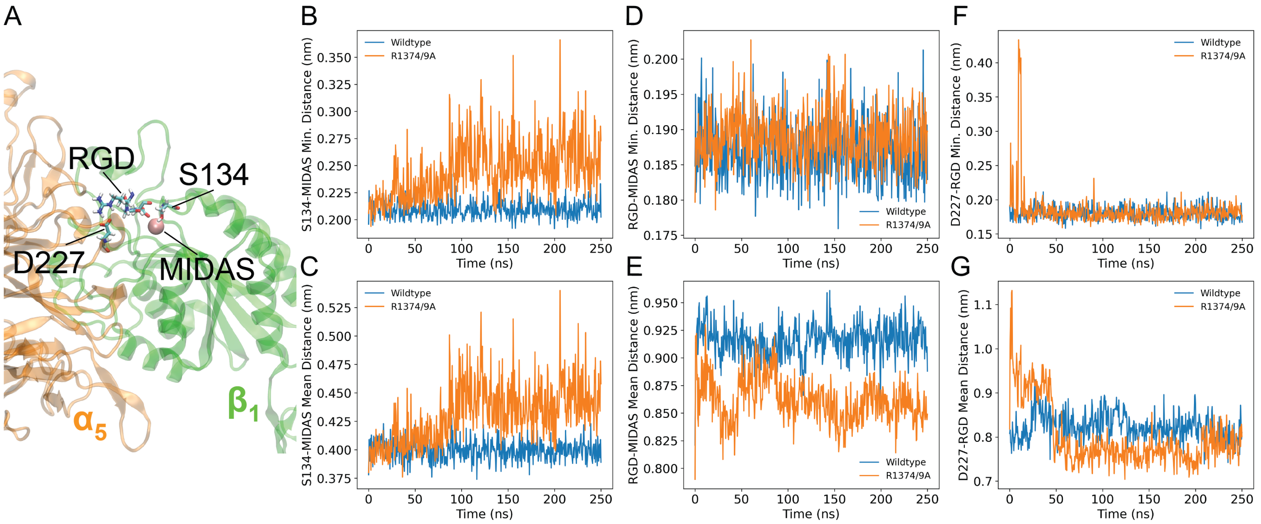


Figure S10: A) RGD site including key molecules such as RGD, S134, and D227, as well as the MIDAS coordination cation. Minimum and mean distances of B-C) S134-MIDAS, D-E) RGD-MIDAS, F-G) D227-RGD.

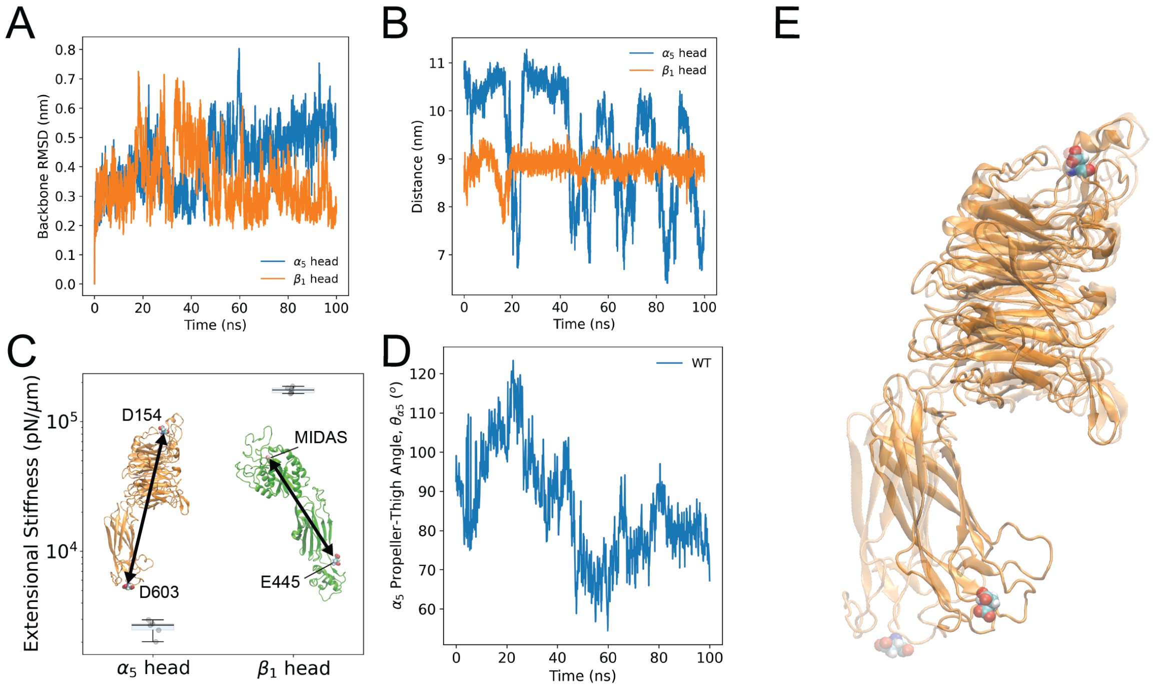


Figure S11: A) Backbone RMSD of independent $\alpha_5$ and $\beta_1$ heads. B) Distance between D603 and D154 ($\alpha_5$) and E445 and MIDAS ($\beta_1$). C) Extensional stiffnesses of $\alpha_5$ and $\beta_1$ as measured by the respective reaction coordinates. D) Propeller-thigh angle on $\alpha_5$. E) First (transparent) and last (opaque) frames of $\alpha_5$ simulation. D603 (top) and D154 (bottom) are shown as references.
